# Multiple ParA/MinD ATPases coordinate the positioning of disparate cargos in a bacterial cell

**DOI:** 10.1101/2022.06.09.495121

**Authors:** Lisa T. Pulianmackal, Jose Miguel I. Limcaoco, Keerthikka Ravi, Sinyu Yang, Jeffrey Zhang, Mimi K. Tran, Matthew J. O’Meara, Anthony G. Vecchiarelli

**Affiliations:** Department of Microbiology and Immunology, University of Michigan, Ann Arbor, MI, 48109, USA; Department of Molecular, Cellular, and Developmental Biology, University of Michigan, Ann Arbor, MI, 48109, USA; Department of Computational Medicine & Bioinformatics, University of Michigan, Ann Arbor, MI, 48109, USA

**Author notes:** **Correspondence** (A.G.V).

## Abstract

In eukaryotes, linear motor proteins govern intracellular transport and organization. In bacteria, where linear motors are absent, the ParA/MinD (A/D) family of ATPases spatially organize an array of genetic- and protein-based cellular cargos. ParA is well known to segregate plasmids and chromosomes, as is MinD for its role in divisome positioning. Less studied is the growing list of ParA/MinD-like ATPases found across prokaryotes and involved in the spatial organization of diverse protein-based organelles, such as <u>B</u>acterial <u>M</u>icrocompartments (BMCs), flagella, chemotaxis clusters, and conjugation machinery. Given the fundamental nature of these processes in both cell survival and pathogenesis, the positioning of these cargos has been independently investigated to varying degrees in several organisms. However, it remains unknown whether multiple A/D ATPases can coexist and coordinate the positioning of such a diverse set of fundamental cargos in the same cell. If so, what are the mechanistic commonalities, variation, and specificity determinants that govern the positioning reaction for each cargo? Here, we find that over a third of sequenced bacteria encode multiple A/D ATPases. Among these bacteria, we identified several human pathogens as well as the experimentally tractable organism, *Halothiobacillus neapolitanus*, which encodes seven A/D ATPases. We directly demonstrate that five of these A/D ATPases are each dedicated to the spatial regulation of a single cellular cargo: the chromosome, the divisome, the carboxysome BMC, the flagellum, and the chemotaxis cluster. We identify putative specificity determinants that allow each A/D ATPase to position its respective cargo. Finally, we show how the deletion of one A/D ATPase can have indirect effects on the inheritance of a cargo actively positioned by another A/D ATPase, stressing the importance of understanding how organelle trafficking, chromosome segregation, and cell division are coordinated in bacterial cells. Together, our data show how multiple A/D ATPases coexist and function to position a diverse set of fundamental cargos in the same bacterial cell. With this knowledge, we anticipate the design of minimal autonomous positioning systems for natural- and synthetic-cargos in bacteria for synthetic biology and biomedical applications.

## Introduction

Actin filaments, microtubules, and the linear motor proteins that walk along them, are well known for spatial organization in eukaryotic cells. In bacteria, however, where linear motor proteins are absent, a widespread family of ParA/MinD (A/D) ATPases spatially organize plasmids, chromosomes, and an array of protein-based organelles, many of which are fundamental to cell survival and pathogenesis. By far the two best studied ATPases, and family namesake, are <u>Par</u>A involved in plasmid <u>par</u>tition and chromosome segregation **(Baxter and Funnell, 2014; Jalal and Le, 2020)**, and MinD involved in divisome positioning **(Lutkenhaus, 2007)**. Less studied is the growing list of A/D ATPases, widespread across prokaryotes, involved in spatially regulating diverse protein-based organelles, such as Bacterial <u>M</u>icro<u>c</u>ompartments (BMCs) **(MacCready et al., 2021; Savage et al., 2010)**, flagella (Schuhmacher et al., 2015b, 2015a), chemotaxis clusters **(Ringgaard et al., 2011; Thompson et al., 2006)**, and conjugation machinery **(Atmakuri et al., 2007)**.

Despite the cargos being so diverse, A/D ATPases share a number of features: (*i*) all form ATP-sandwich dimers **(Shan, 2016)**, (*ii*) dimerization forms an interface for binding a positioning matrix - the nucleoid for ParA-like ATPases **(Hester and Lutkenhaus, 2007; Kiekebusch et al., 2012)** or the inner membrane for MinD-like ATPases **(Hu and Lutkenhaus, 2003; Szeto et al., 2002)**, and (*iii*) dimerization also forms a binding site for a cognate partner protein that connects an ATPase to its cargo and stimulates its release from the positioning matrix. For example, in chromosome segregation, the ParA partner is ParB, which loads onto a centromere-like site, called *parS*, to form a massive complex on the chromosome near the origin of replication (*OriC*) **(Jalal and Le, 2020)**. This ParB-*parS* complex locally stimulates ParA ATPase activity and nucleoid release, which generates ParA gradients on the nucleoid. Segregation ensues as sister chromosomes chase nucleoid-bound ParA gradients in opposite directions **(Vecchiarelli et al., 2010)**. Therefore, unlike the mitotic-spindle apparatus used in eukaryotic chromosome segregation, prokaryotes use a fundamentally different mode of spatial organization - A/D ATPases make waves on biological surfaces to position their respective cargos.

Chromosome segregation, cell division positioning, and organelle trafficking reactions have been independently investigated to varying degrees in several prokaryotes. Yet, it remains unknown how many A/D ATPases can be encoded in a single bacterium to position multiple disparate cargos, or how bacteria spatiotemporally coordinate the positioning of such a diverse set of fundamental cargos in the same cell. Here, we find that a third of sequenced bacteria encode for multiple A/D ATPases. Among these bacteria, we identified several human pathogens as well as the non-pathogenic and experimentally tractable organism, *Halothiobacillus neapolitanus* (*H. neapolitanus* hereafter), with seven putative A/D ATPases. Neighborhood analysis of the A/D ATPase genes in *H. neapolitanus* implicate several putative cargos, six of which are already known to be positioned by an A/D ATPase in other bacteria. The tractability and number of A/D ATPases make *H. neapolitanus* a valuable tool for investigating how bacteria coordinate the processes of chromosome segregation and cell division with organelle trafficking – a well-studied question in eukaryotic cells that remains unaddressed in prokaryotes.

Additionally, the mechanistic variations and specificity determinants that govern the positioning of such a diverse set of cellular cargo also remain unclear. This is because A/D-based positioning reactions are typically studied independently of one another and in model bacteria with few A/D ATPases. Here we use genetics and cell biology to assign five of the A/D ATPases in *H. neapolitanus* to their cargos. Our findings show that each ATPase is directly dedicated to the positioning and faithful inheritance of a specific cargo type. We then show how the deletion of one A/D ATPase can have indirect effects on the inheritance of disparate cargos positioned by other A/D ATPases via defects in DNA replication, chromosome segregation, and/or cell division. We also, for the first time, provide evidence that flagella positioning influences the spatial regulation of chemotaxis clusters. Finally, we identify putative sequence- and structural-determinants that uniquely link each A/D ATPase to a specific cargo, ultimately allowing these related ATPases to coexist and function in the same cell. Together, our study probes mechanistic commonality and variation in the most widespread ATPase family used in the spatial regulation of diverse cellular cargos across prokaryotes, and all within a single cell.

## Results

### A third of sequenced bacteria encode for multiple ParA/MinD family ATPases

It is unclear how many ParA/MinD (A/D) family ATPases can be encoded in a single organism to position multiple disparate cargos in the same cell. To answer this question, we performed an extensive tBLASTn analysis using a consensus protein sequence, generated from well-studied A/D ATPases, as the query (see methods). As already established **(Koonin, 1993)**, we found that ∼ 95% of bacteria from the NCBI Reference Sequence (RefSeq) database encode for at least one A/D ATPase **(Table 1)**. These hits were binned by bacterial species, which were then ordered by the number of A/D hits. From this initial list, we found many bacterial genomes encoding 10 to 20 A/D ATPases. However, these bacteria with the most A/D ATPases had their genomes encoded on multiple plasmids and chromosomes, each of which encode its own ParA-based DNA segregation system **(Tilly et al., 2012)**. For this study, we were specifically focused on understanding how multiple A/D ATPases coexist and coordinate the positioning of *disparate* cargos in the same cell. Therefore, we further filtered our dataset to identify bacteria encoding multiple A/D ATPases, but only one chromosome and no stable plasmids **(Table S1)**. Even after accounting for bacteria with genomes encoded on multiple genetic elements, our bioinformatic analysis revealed that more than a third of sequenced bacteria encode multiple A/D ATPases **(Figure 1A, Table 1)**.

**Figure 1:**
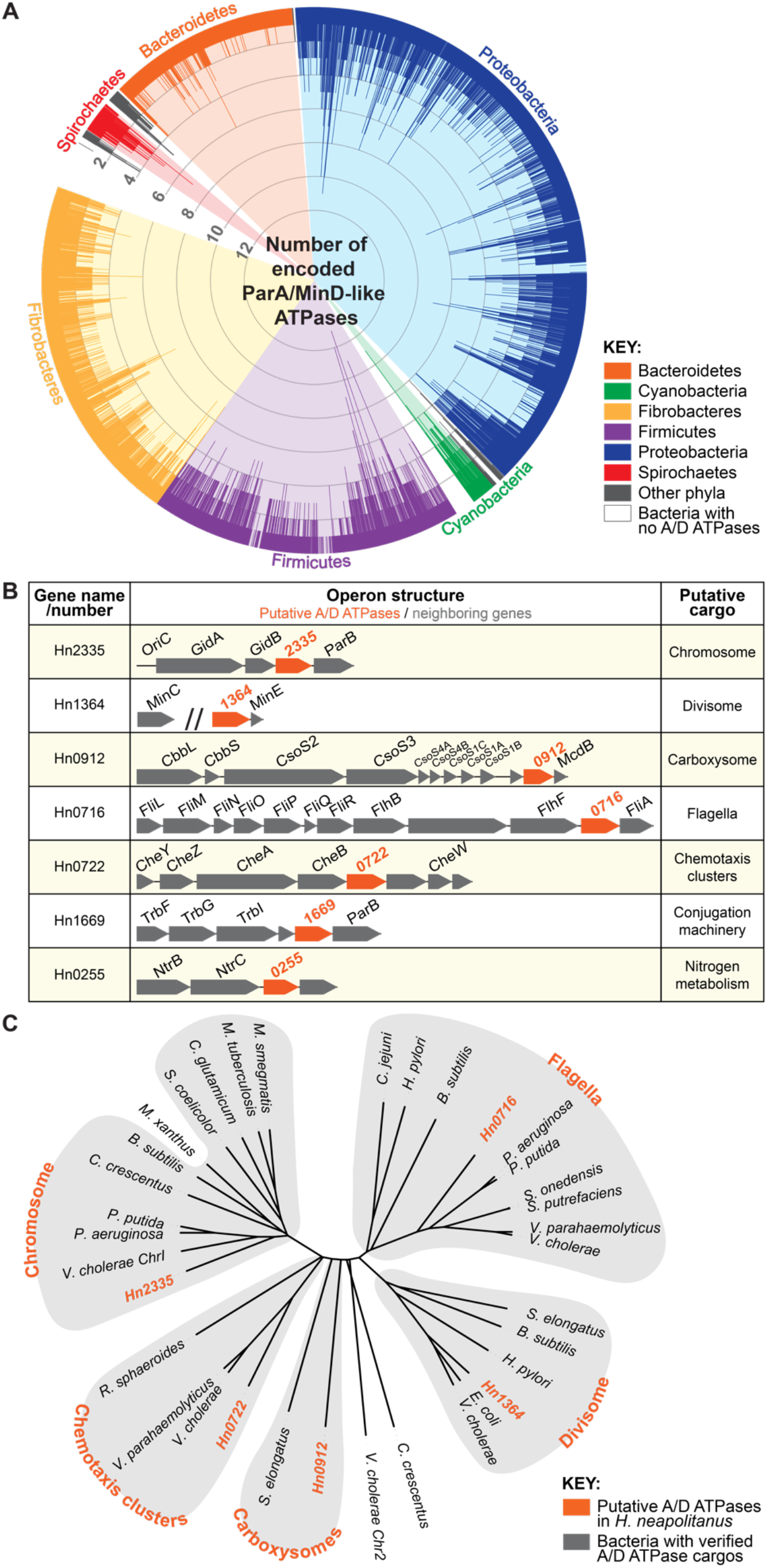
ParA/MinD-like ATPases are widespread in bacteria. **(A)** 95% of sequenced bacterial genomes encode for at least one A/D ATPase and more than a third encode for multiple. Each spike represents a bacterial species and spike length indicates the number of unique A/D hits per bacterium. **(B)** *H. neapolitanus* encodes for seven putative ParA/MinD-like positioning systems. GNA implicates the putative cargos associated with each putative A/D ATPase. **(C)** MSA of each A/D ATPase from *H. neapolitanus* against experimentally-verified ParA/MinD family members further implicates the putative cargos identified by GNA. Each of the putative A/D ATPases in *H. neapolitanus* cluster with family members shown to position the indicated cellular cargos (orange) in other bacteria.

### Gene neighborhood analysis of A/D ATPases in *H. neapolitanus* implicate cellular cargos

We next set out to determine how multiple A/D ATPases can coexist in the same cell to position disparate cargos. To address this question, we identified an organism from our list of bacteria encoding multiple A/D ATPases **(Table 1)**. Among the top 1% of bacteria (encoding six or more A/D ATPases), we identified several human pathogens as well as the non-pathogenic and experimentally tractable organism, *H. neapolitanus* - a slow-growing, sulfur-oxidizing chemoautotroph that encodes seven putative A/D ATPases on one chromosome **(Figure 1B)**. Spatial regulation by A/D ATPases has largely been studied in fast-growing model bacteria. We intentionally chose a slow growing bacterium (6 hr doubling time) because the infrequent DNA segregation and cell division events of *H. neapolitanus* allowed for larger observation windows and a more direct view into the dynamics of organelle trafficking. The tractability, slow-growth rate, and abundance of A/D ATPases made *H. neapolitanus* an ideal choice for further study.

To determine whether the A/D ATPase hits in *H. neapolitanus* were indeed spatial regulators, we performed a gene neighborhood analysis (GNA) to identify putative cargos **(Figure 1B)**. GNA allows us to infer function because A/D ATPase genes are often encoded near cargo-associated loci. For example, the ParAB*S* chromosome segregation system is typically encoded near *OriC* **(Livny et al., 2007)**. Strikingly, the putative cargos we identified using this approach includes the chromosome and all known protein-based cargos of the A/D ATPase family (Lutkenhaus, 2012; Vecchiarelli et al., 2012). While spatial regulation of these diverse cargos has been individually studied in many different model bacteria, their coordinated positioning by multiple A/D ATPases has never been investigated together in one organism.

As a second line of bioinformatic evidence linking each A/D ATPase to a specific cargo, we investigated the conservation of the A/D ATPase gene neighborhoods using FlaGs (Flanking Genes) analysis **(Saha et al., 2021)**. FlaGs analysis predicts functional associations by taking a list of NCBI protein accessions as input and clusters neighborhood-encoded proteins into homologous groups. Homologs of each A/D ATPase in *H. neapolitanus* were identified using BLASTp, and top hits were used as input for FlaGs analysis **(Figure S1)**. The analysis shows strong conservation of the A/D ATPase gene neighborhoods across multiple bacterial phyla, further implicating the putative cargos. Due to the limited data on A/D ATPases associated with conjugation **(Atmakuri et al., 2007)** or nitrogen metabolism, we excluded these hits from further investigation in this study.

With the remaining five A/D ATPases, we performed a multiple sequence alignment against ParA/MinD family members that have been previously established to position a cellular cargo **(Figure 1C)**. Conveniently, each A/D ATPase in *H. neapolitanus* clustered with a specific family member known to position a specific cellular cargo in other bacteria - the chromosome (ParA **(Jalal and Le, 2020)**), the divisome (MinD **(Raskin and De Boer, 1999)**), the carboxysome (McdA **(MacCready et al., 2021; Savage et al., 2010)**), the flagellum (FlhG **(Schuhmacher et al., 2015b, 2015a)**) and the chemotaxis cluster (ParC **(Ringgaard et al., 2011, 2014; Thompson et al., 2006)**). These data provide a third line of evidence that further implicates the putative cargos identified by our GNA. We next sought to directly identify the role of each A/D ATPase in positioning the cargos implicated by bioinformatics.

### The ParAB system is required for chromosome segregation in *H. neapolitanus*

Chromosome segregation prior to cell division is critical for cellular survival. In most bacteria, faithful chromosome segregation and inheritance are mediated by a ParAB system encoded near *OriC* **(Livny et al., 2007)**. Without ParAB, DNA is asymmetrically inherited, resulting in anucleate and polyploid cells, and reduced cell fitness or death **(Jalal and Le, 2020)**. In *H. neapolitanus*, there is a putative *parAB* system encoded near *OriC* **(Figure 2A)**. To image chromosome segregation, the ParB homolog encoded downstream of the putative *parA* gene (*Hn2335*) was fused to the fluorescent protein <u>m</u>onomeric <u>N</u>eon <u>G</u>reen (mNG). ParB-mNG was observed as one or two puncta per cell **(Figure 2B, left)**. Population analysis showed that shorter cells had a single focus at mid-cell, whereas longer cells had two foci at the quarter positions of the cell **(Figure 2B, left)**. When the putative *parA* gene (*Hn2335*) was deleted, ParB-mNG foci were randomly positioned regardless of cell length **(Figure 2C)**, or completely absent in 25% of cells **(Figure S2D)**. When present, ParB foci were also significantly brighter compared to that of WT (wild-type) cells **(Figure S2E)**. The data suggest that replicated chromosomes were no longer faithfully segregated, resulting in anucleate and polyploid cells.

**Figure 2:**
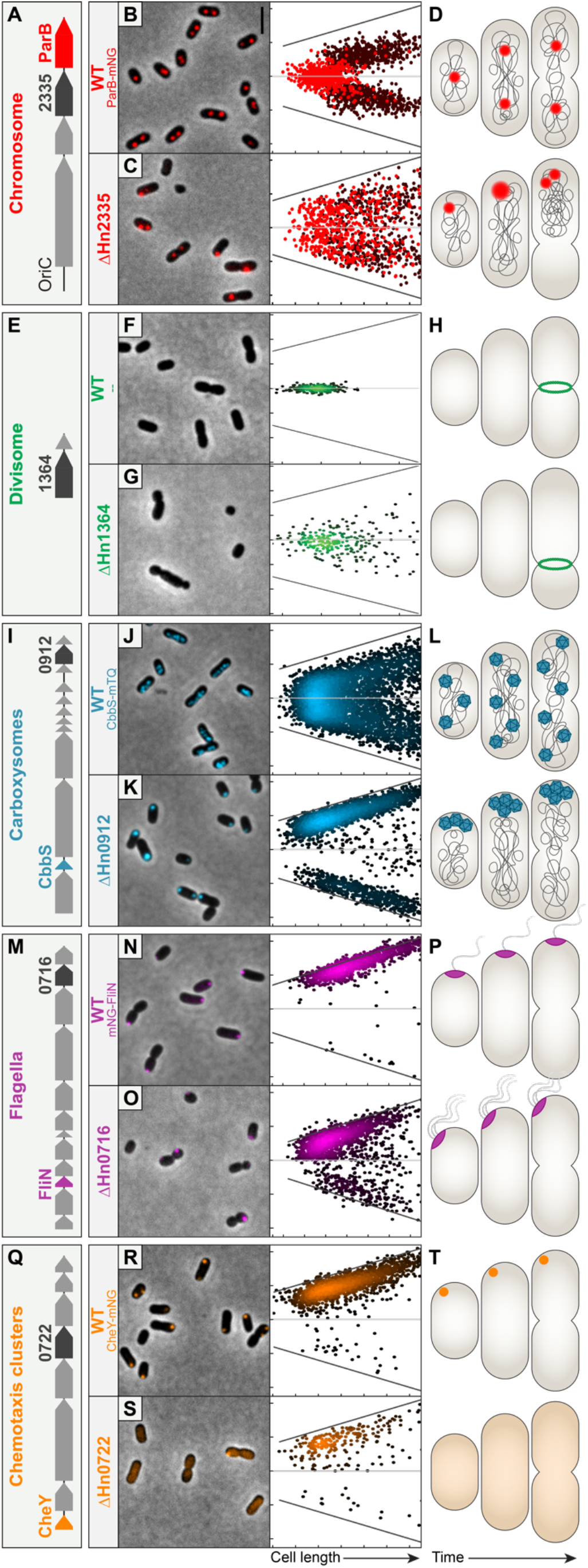
Each A/D ATPase in *H. neapolitanus* positions a specific cargo. (A-D) *Hn2335* is required for chromosome segregation. **(A)** *Hn2335* is found near *OriC* and has a ParB-homolog encoded downstream. **(B, C)** The origin region of the chromosome was tagged by labelling the ParB homolog. Light red: 1 focus/cell; dark red: 2 foci/cell. Short WT cells had a single focus at mid-cell, whereas longer cells had two foci at the quarter positions. Δ*Hn2335* cells displayed random positioning of ParB foci regardless of cell length. **(D)** Cartoon diagrams depict chromosome segregation in WT and Δ*Hn2335* cells. **(E-H) *Hn1364* is required for divisome positioning. (E)** *Hn1364* is found upstream of *minE*. **(F, G)** Divisome positioning was determined by the location of constriction sites. Each dot on the graph represents one identified constriction site. In WT cells, constriction sites were at mid-cell. In Δ*Hn1364*, constriction sites were found across the cell length. **(H)** Cartoon diagrams depict cell division in WT and Δ*Hn1364* cells. **(I-L) Carboxysome positioning is determined by McdA. (I)** *Hn0912* is found near genes encoding carboxysome shell proteins and RuBisCO. **(J, K)** Carboxysomes were visualized by labelling the small subunit of the Rubisco enzyme, *cbbS*. In WT cells, carboxysomes are distributed along the cell length. In Δ*mcdA*, carboxysomes formed large polar foci at one or both poles. **(L)** Cartoon diagrams depict carboxysome distribution in WT and Δ*Hn0912* cells. **(M-P) *Hn0716* is required for regulating flagella position and copy number. (M)** *Hn0716* is found near flagella-associated genes. **(N, O)** Flagella localization was visualized by labelling a component of the flagellar basal body, *fliN*. WT cells had a single polar FliN focus. In Δ*Hn0716* cells, FliN foci were more randomly positioned along the cell length. **(P)** Cartoon diagrams depict flagella localization and number in WT and Δ*Hn0716* cells. **(Q-T) *Hn0722* is required for chemotaxis cluster positioning. (Q)** *Hn0722* is found near chemotaxis-associated genes. **(R, S)** Chemotaxis clusters were visualized by labelling the response regulator, *cheY*. WT cells had a single CheY polar focus. Δ*Hn0722* mutant cells typically had no CheY foci. **(T)** Cartoon diagrams depict chemotaxis clusters in WT and Δ*Hn0722* cells. **(All images)** Scale bar: 2 μm. **(All graphs)** Cells were analyzed and quantified using MicrobeJ. On the x-axis, cells are sorted by increasing cell length. The y-axis represents the distance from mid-cell in microns; the center horizontal line equates to a distance of zero from mid-cell. Each dot represents where a focus was found along the length of the cell. For flagella, chemotaxis, and carboxysome graphs, the cell pole closest to a focus was oriented at the top; foci in the bottom half of the graph indicate the presence of a second focus. Graph axes for chromosome, carboxysome, flagella, and chemotaxis-labelled mutants: x-axis range (cell length): 0.8 to 2.1 μm; y-axis range (distance from mid-cell): -1.1 to 1.1 μm. X-axis of Δ*flhG* cells is 0.5 to 1.8 μm. Graph axes for divisome: x-axis range (cell length): 1.5 to 3.0 μm; y-axis range (distance from mid-cell): -1.5 to 1.5 μm.

We then performed time-lapse microscopy of ParB-mNG foci and SYTOX-stained nucleoids to observe chromosome segregation in real time. Newborn WT cells have a single ParB focus at mid-cell, which then splits into two foci that bidirectionally segregate towards the quarter positions of the growing cell **(Figure S2F and Movie 1A)**. Foci positioning at the cell quarters was maintained, which then became the mid-cell position of each daughter cell following division. Faithful chromosome segregation and inheritance were lost when the putative *parA* gene (*Hn2335*) was deleted **(Figure S2G and Movies 1B)**. Many cells had a single large ParB focus at a cell pole that did not split and all DNA was concentrated into a single daughter cell upon division. These polyploid daughters continued to divide, while the anucleate daughters were no longer viable. These data **confirmed that the increased** ParB foci intensities in the deletion mutant represents an increase in chromosome copy number in these cells. In summary, the A/D ATPase encoded by *Hn2335* is required for faithful chromosome segregation **(Figure 2D)** and we will henceforth refer to this protein as ParA.

### The MinCDE system aligns cell division at mid-cell in *H. neapolitanus*

Proper positioning of the divisome ensures that when a cell divides, both daughter cells are roughly equal in length. Without the Min system, division occurs at any nucleoid-free region **(Yu and Margolin, 1999)**, producing anucleate <u>mini</u>cells, the products of polar divisions. Our bioinformatic analyses showed that the protein encoded by *Hn1364* is a MinD homolog, and immediately downstream of this gene in the same operon is a gene encoding a MinE homolog **(Figure 2E)**. To determine if *Hn1364* was indeed involved in divisome positioning, we analyzed dividing cells and identified their constriction sites relative to cell length **(Figure 2F-G)**. WT cells divided close to mid-cell **(Figure 2F and Figure S3D)** while the Δ*Hn1364* cells divided asymmetrically **(Figure 2G and Figure S3E)**. Population analysis of dividing cells identified mid-cell constriction sites in 97% of WT cells compared to only 41% of Δ*Hn1364* cells **(Figure S3F and Movie 2)**. In addition to single asymmetric division events, 9% of dividing cells in the Δ*Hn1364* mutant population formed multiple division sites simultaneously along the cell length **(Figure S3G and Movie 2C)**. The unequal divisions resulted in greater variation in cell length **(Figure S3H)**. Overall, our findings show that *Hn1364* is critical for positioning the divisome at mid-cell **(Figure 2H)** and we will henceforth refer to this protein as MinD.

### The A/D ATPase, McdA, encoded in the carboxysome operon positions carboxysomes

Bacterial microcompartments, or BMCs, are large icosahedral protein-based organelles that encapsulate sensitive metabolic reactions to provide prokaryotes with distinct growth advantages **(Kerfeld et al., 2018)**. Despite their importance, little is known about how BMCs are spatially regulated. The model BMC is the carbon-fixing carboxysome found in cyanobacteria and proteobacteria **(Turmo et al., 2017)**. It was recently found that an A/D ATPase, we termed <u>M</u>aintenance of <u>c</u>arboxysome <u>d</u>istribution protein A (McdA), is widespread among carboxysome-containing bacteria, including *H. neapolitanus* **(Figure 2I) (Maccready and Vecchiarelli, 2021)**. McdA spaces carboxysomes on the nucleoid along with its partner protein, McdB **(MacCready et al., 2018, 2021)**. To demonstrate its requirement for carboxysome positioning, we visualized carboxysomes by labelling the small subunit of the encapsulated Rubisco enzyme (*cbbS*) with mTurquoise2 to form CbbS-mTQ. As previously shown, in WT cells, carboxysomes are distributed down the cell length **(Figure 2J)**. In the deletion mutant (Δ*Hn0912*), carboxysome aggregates form a large polar focus at one or occasionally both poles **(Figure 2K)**. These data are consistent with our previous observations using TEM, which showed that foci in the mutant population represent an aggregation of assembled carboxysomes **(MacCready et al., 2021)**.

We extend our previous findings here with long-term time-lapse microscopy. In WT cells, carboxysomes are dynamically positioned along the cell length across multiple generations **(Figure S4F and Movie 3A)**. In the deletion mutant, carboxysome aggregates were stagnant **(Figure S4G and Movie 3B)**. Together, our findings show that the A/D ATPase encoded within the carboxysome operon, we termed McdA, is essential for distributing carboxysomes across the cell length and ensuring organelle homeostasis following division **(Figure 2L)**.

### *Hn0716* is required for regulating flagella number and positioning

Flagella are external filamentous structures that allow for bacterial motility. Bacteria vary in flagella location, number, and pattern. Many bacteria encode for an A/D ATPase called FlhG in their flagellar operon, which is essential for diverse flagellation patterns in many bacteria **(Schuhmacher et al., 2015b)**, yet the mechanisms remain unclear. Deletion of *flhG* typically results in changes in flagella number, location, and motility. Our bioinformatic analysis suggests that *Hn0716* within the flagella operon encodes an FlhG homolog **(see Figure 1C)**. To determine if *Hn0716* is involved in flagellar spatial regulation, we first visualized a mNG fusion of FliN, which is a component of the flagellar basal body that assembles at the cytoplasmic face of the membrane **(Figure 2M)** (Chang et al., 2020). WT cells had a single mNG-FliN focus at the extreme cell pole **(Figure 2N)**. In Δ*Hn0716* cells, FliN foci were no longer faithfully positioned **(Figure 2O)** and cells were more likely to have multiple foci **(Figure S5D)**. The foci in Δ*Hn0716* cells were also lower in fluorescence intensity compared to WT **(Figure S5E)**. These data suggest that *Hn0716* is required for positioning flagellar machinery at a single pole in *H. neapolitanus*.

We next set out to determine whether these alterations to FliN positioning affected cell motility. Motility assays found that Δ*Hn0716* cells were not motile **(Figure S5F)**. Loss of motility could be due to flagellar mispositioning, a loss of flagella, or hyper-flagellation. To image flagella, we engineered flagellin^T185C^, which allows for fluorescent labeling of flagella via the addition of a cysteine-reactive maleimide stain to the media **(Kühn et al., 2018)**. We found that WT *H. neapolitanus* cells are monotrichous, with a single polar flagellum emanating from the FliN focus **(Figure S5G)**. Δ*Hn0716* cells also had flagella emanating from FliN foci, however, the cells were hyper-flagellated, with flagella often bundled together as tufts. We conclude that *Hn0716* is required for regulating flagella number and position in *H. neapolitanus* **(Figure 2P)** and we will henceforth refer to this protein as FlhG.

### *Hn0722* is required for chemotaxis cluster assembly and positioning

Directing bacterial motility are large hexagonal arrays called chemotaxis clusters, comprised of chemoreceptors, an adaptor protein (CheW), and a kinase (CheA). Several mechanisms have evolved to control both the number and positioning of chemotaxis clusters, including the use of A/D ATPases (called ParC in *Vibrio* species or PpfA in *R. sphaeroides*) **(Ringgaard et al., 2011; Roberts et al., 2012)**. Deletion of the A/D ATPase alters chemotaxis cluster number and positioning in cells, which results in a reduction in swarming motility. Our bioinformatics analysis showed that the protein encoded by *Hn0722* is a ParC homolog within the chemotaxis operon of *H. neapolitanus* **(see Figure 1C)**. To determine if *Hn0722* is important for spatially regulating chemotaxis clusters, we first imaged chemotaxis clusters by fusing mNG to CheY **(Figure 2Q)**. CheY is a response regulator that is phosphorylated by CheA, and has previously been shown to colocalize with chemotaxis clusters in *E. coli* **(Kentner and Sourjik, 2009; Sourjik and Berg, 2000)**. Consistent with electron micrographs of chemotaxis clusters in *H. neapolitanus* (Briegel et al., 2009), CheY-mNG formed a single focus immediately proximal to one cell pole in ∼ 85% of WT cells **(Figure 2R, Figure S6D)**. However, when *Hn0722* was deleted, the CheY-mNG signal was diffuse in ∼ 80% of the cell population **(Figure 2S, Figure S6D)**. In the ∼ 20% of Δ*Hn0722* cells that had a CheY-mNG focus, the foci were significantly lower in intensity **(Figure S6E)**. Together, we conclude that *Hn0722* is required for chemotaxis cluster assembly and positioning in *H. neapolitanus* **(Figure 2T)** and we will henceforth refer to this protein as ParC.

### Cargo positioning is not directly controlled by A/D ATPases encoded at distant loci

We have thus far provided direct evidence showing that five A/D ATPases position five disparate cellular cargos in *H. neapolitanus* **(Figure 2)**. We next asked if each of the five positioning reactions occurred independently from each other, or whether they exhibited cross-talk. To answer this question, we individually deleted each A/D ATPase in every cargo-labeled background strain **(Figure 3)**. We found carboxysome **(Figure 3C)** and flagella **(Figure 3D)** positioning were largely unaffected by the deletion of distant A/D ATPases. Intriguingly, chromosome **(Figure 3A)**, divisome **(Figure 3B)**, and chemotaxis cluster **(Figure 3E)** positioning were all influenced in Δ*flhG* cells, albeit with intermediate phenotypes when compared to deleting the dedicated A/D ATPase **(Figure 3, bold boxes)**. Chemotaxis cluster positioning was also mildly influenced in Δ*parA* cells **(Figure 3E)**. Together our data show that each A/D ATPase is dedicated to the positioning of a specific cargo type, and not *directly* involved in the positioning of other cargos. However, our data at the cell-population level also unveiled potential coordination, crosstalk, and/or interdependencies among certain positioning reactions. In the next three sections, we dissect how the deletion of one A/D ATPase can indirectly affect the positioning and inheritance of cellular cargos positioned by another A/D ATPase.

**Figure 3:**
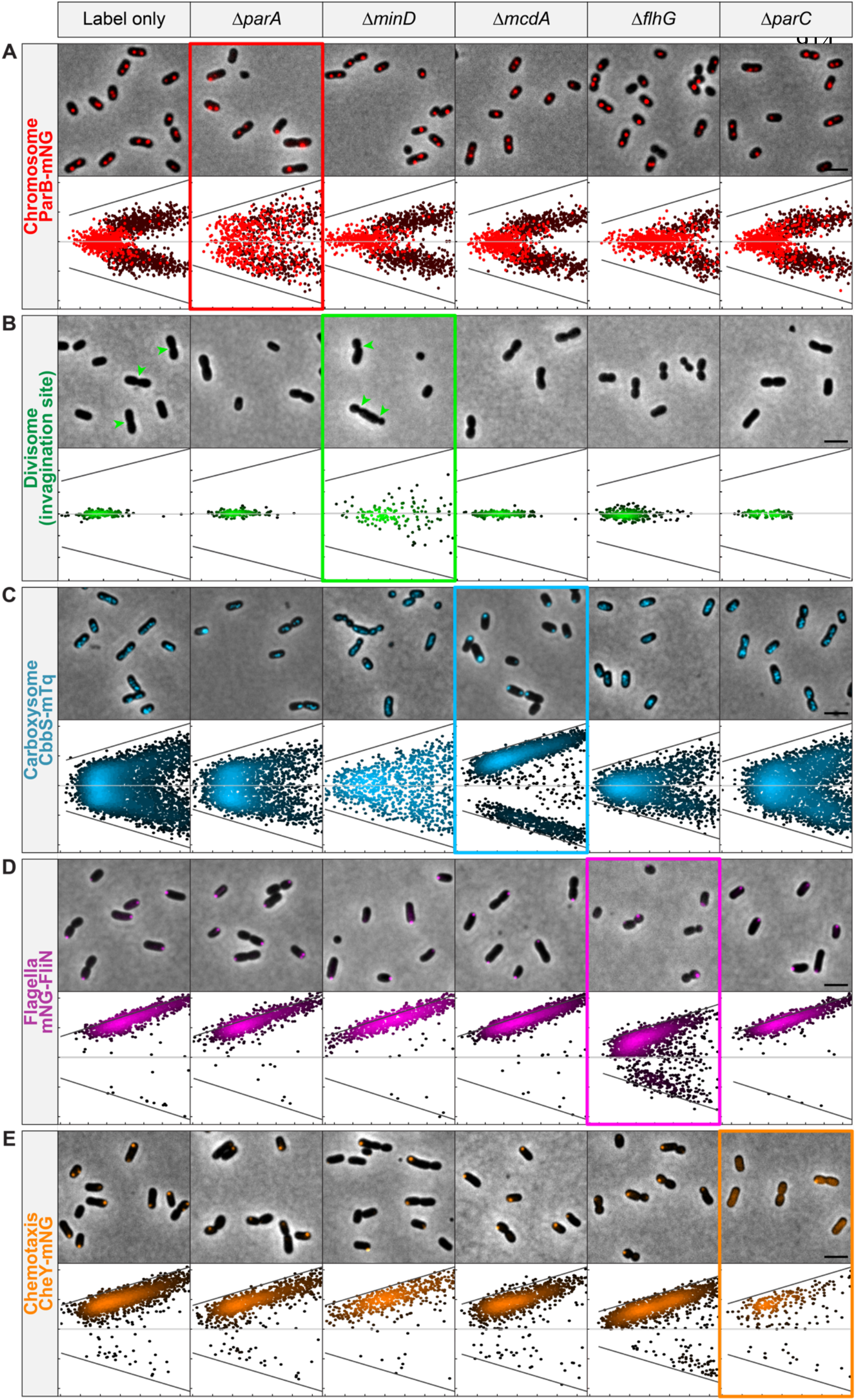
Cargo positioning is not directly controlled by A/D ATPases encoded at distant loci. Each of the five cellular cargos were fluorescently tagged as indicated (Left): **(A)** chromosome (ParB-mNG), **(B)** divisome (invagination site), **(C)** carboxysomes (CbbS-mTQ), **(D)** flagella (mNG-FliN), and **(E)** chemotaxis clusters (CheY-mNG). The “Label Only” column shows the WT positioning of each of the fluorescent cargos. Deletion of an A/D ATPase resulted in the mislocalization of only its specific cargo (bolded rectangles). Graph axes for chromosome, carboxysome, flagella, and chemotaxis-labelled mutants: x-axis range (cell length): 0.8 to 2.1 μm; y-axis range (distance from mid-cell): -1.1 to 1.1 μm. All x-axes of Δ*flhG* cells are 0.5 to 1.8 μm. Graph axes for divisome: x-axis range (cell length): 1.5 to 3.0 μm; y-axis range (distance from mid-cell): -1.5 to 1.5 μm.

### Deletion of *parA, minD*, or *flhG* results in anucleate cells via three different mechanisms

We have shown how the deletion of *parA* results in a significant fraction of anucleate cells because ParA is directly required for chromosome segregation and inheritance following cell division **(Figure 4A)**. Deleting *minD* or *flhG* also resulted in anucleate cells, but the mechanism was different in each case. In Δ*minD* cells, chromosome positioning **(Figure 4B)** and ParB foci intensities **(Figure 4C)** were similar to that of WT, showing that chromosome segregation was still active. Instead, it was divisome mispositioning in Δ*minD* cells and subsequent asymmetric cell division **(Figure 4D)** that indirectly caused asymmetric chromosome inheritance and anucleate cell formation.

**Figure 4:**
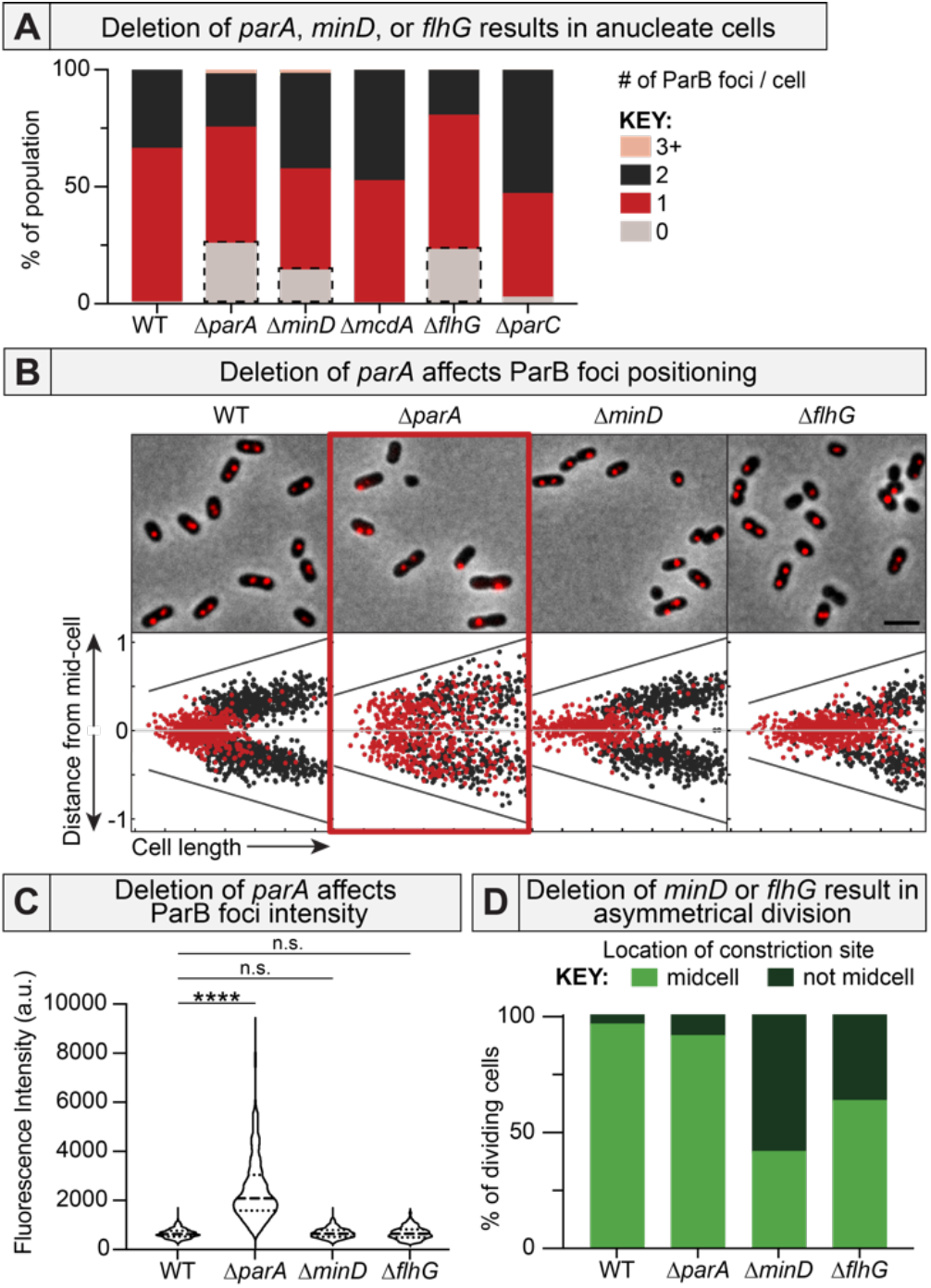
Anucleate cells form via three different mechanisms. **(A)** Deletion of the A/D ATPase required for chromosome, divisome, or flagellar positioning resulted in anucleate cells (grey boxes). **(B)** Only *ΔparA* cells had mispositioned of ParB foci. Cells with a single ParB focus are red. Cells with two ParB foci are black. Δ*flhG* cells maintained a single ParB focus in long cells, suggesting a DNA replication defect. **(C)** ParB foci are brighter only in Δ*parA* cells. **(D)** Deletion of *minD* or *flhG* resulted in divisome mispositioning. Constriction sites were considered “mid-cell” when found within 5% of the cell center along the long axis.

In Δ*flhG* cells, chromosome positioning **(Figure 4B)** and ParB foci intensities **(Figure 4C)** were also similar to that of WT, again suggesting that chromosome segregation was unaffected. But, divisome mispositioning was not as severe as Δ*minD* cells **(Figure 4D, Figure 3B)**, suggesting that anucleate cells were forming via a third distinct mechanism. Intriguingly, Δ*flhG* cells were less likely to have two ParB foci compared to WT cells **(Figure 4A)**. Instead, pre-divisional cells still had a single ParB focus **(Figure 4B)** with intensities that suggested the presence of only a single chromosome copy **(Figure 4C)**. Time-lapse microscopy confirmed that ParB foci positioning was actively maintained **(Movie 4)**. Therefore, anucleate Δ*flhG* cells are likely formed due to defects in chromosome replication and/or premature cell division; a question of future study.

Together, the findings emphasize the importance of probing the functional relationships of A/D ATPases with each other and the bacterial cell cycle, when occupying the same organism.

### Anucleate cells inherit carboxysomes

McdA uses the nucleoid as a positioning matrix for distributing carboxysomes **(MacCready et al., 2018, 2021)**. Therefore, it was surprising that in the Δ*parA* mutant population, anucleate cells retained carboxysomes **(Figure 5A)**. To determine if anucleate cells synthesized carboxysomes *de novo* or carboxysomes were somehow still inherited following division, we performed time-lapse microscopy on Δ*parA* cells with fluorescent carboxysomes **(Figure 5B and Movie 5A)**. Intriguingly, anucleate cells indeed inherited carboxysomes, but in the most unexpected fashion. In dividing Δ*parA* cells that failed to split their ParB-mNG foci, carboxysomes in the to-be-anucleate cell bundled up immediately adjacent to the division plane. Carboxysome bundling was coincident with the chromosome spooling action that occurred at the invaginating septum just prior to complete division and asymmetric chromosome inheritance **(see Figure S2G and Movie 1B)**. After septation was complete, the massive carboxysome bundle was explosively liberated from the new pole of the anucleate cell, resulting in multiple free-diffusing carboxysome foci **(Figure 5B and Movie 5A)**. Anucleate cells harboring carboxysomes did not divide further. We found that anucleate cells in the Δ*minD* **(Figure 5C and Movie 5B)** and Δ*flhG* **(Figure 5D and Movie 5C)** cell populations also inherited carboxysomes via the same mechanism. Together the data show how carboxysome trafficking and distribution by McdA are dependent upon faithful chromosome segregation. However, anucleate cells can still inherit carboxysomes in *parA, minD*, or *flhG* deletion strains because carboxysomes are scraped off of missegregated chromosomes that are spooled through the invagination site during septation. We speculate that all mesoscale cargos using asymmetrically inherited nucleoids as a positioning matrix would show the same mode of inheritance.

**Figure 5:**
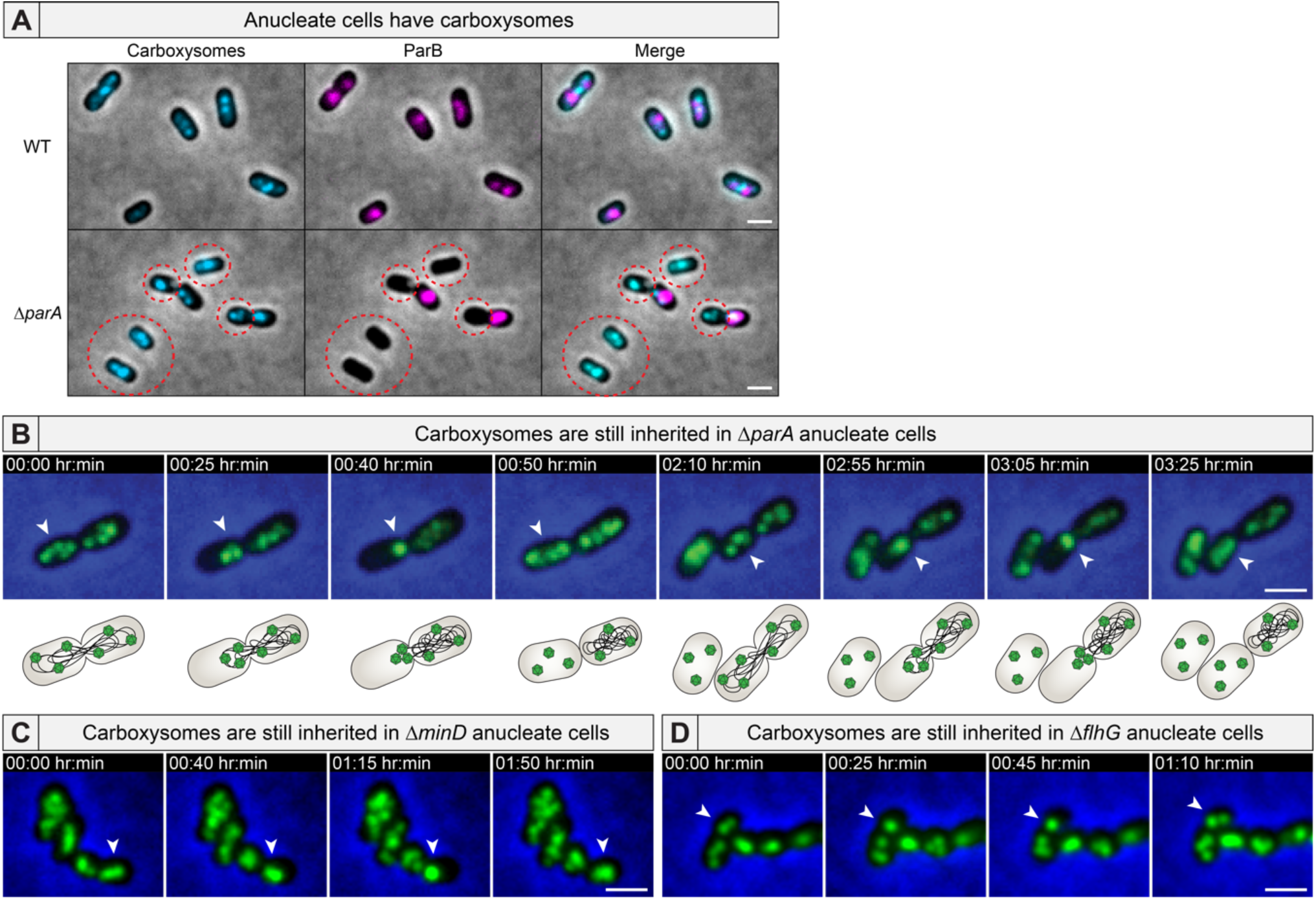
Anucleate cells inherit carboxysomes. **(A)** Carboxysomes are present in anucleate cells (dashed circle). **(B)** Time-lapse microscopy shows that carboxysomes in Δ*parA* cells are bound to the nucleoid, but are inherited in anucleate cells via release from the extruded chromosome. Carboxysomes are also inherited in **(C)** Δ*minD* anucleate cells and **(D)** Δ*flhG* anucleate cells via the same mechanism. **(All videos)** White arrows highlight carboxysome bundling. Scale bar: 1 μm

### Deletion of *parA, minD* or *flhG* influence chemotaxis cluster assembly

The A/D ATPase that positions chemotaxis clusters in *Rhodobacter sphaeroides* uses the nucleoid as its positioning matrix **(Roberts et al., 2012)**. Therefore, we suspected that A/D ATPase deletions resulting in anucleate cells would indirectly impact the spatial regulation of chemotaxis clusters in these strains. Indeed, we found that *parA, minD*, and *flhG* deletion strains, all of which form anucleate cells **(see Figure 4A)**, had a corresponding increase in cells lacking chemotaxis clusters **(Figure 6A)**, and when cells had foci, they were notably dimmer compared to WT **(Figure 6B)**.

**Figure 6:**
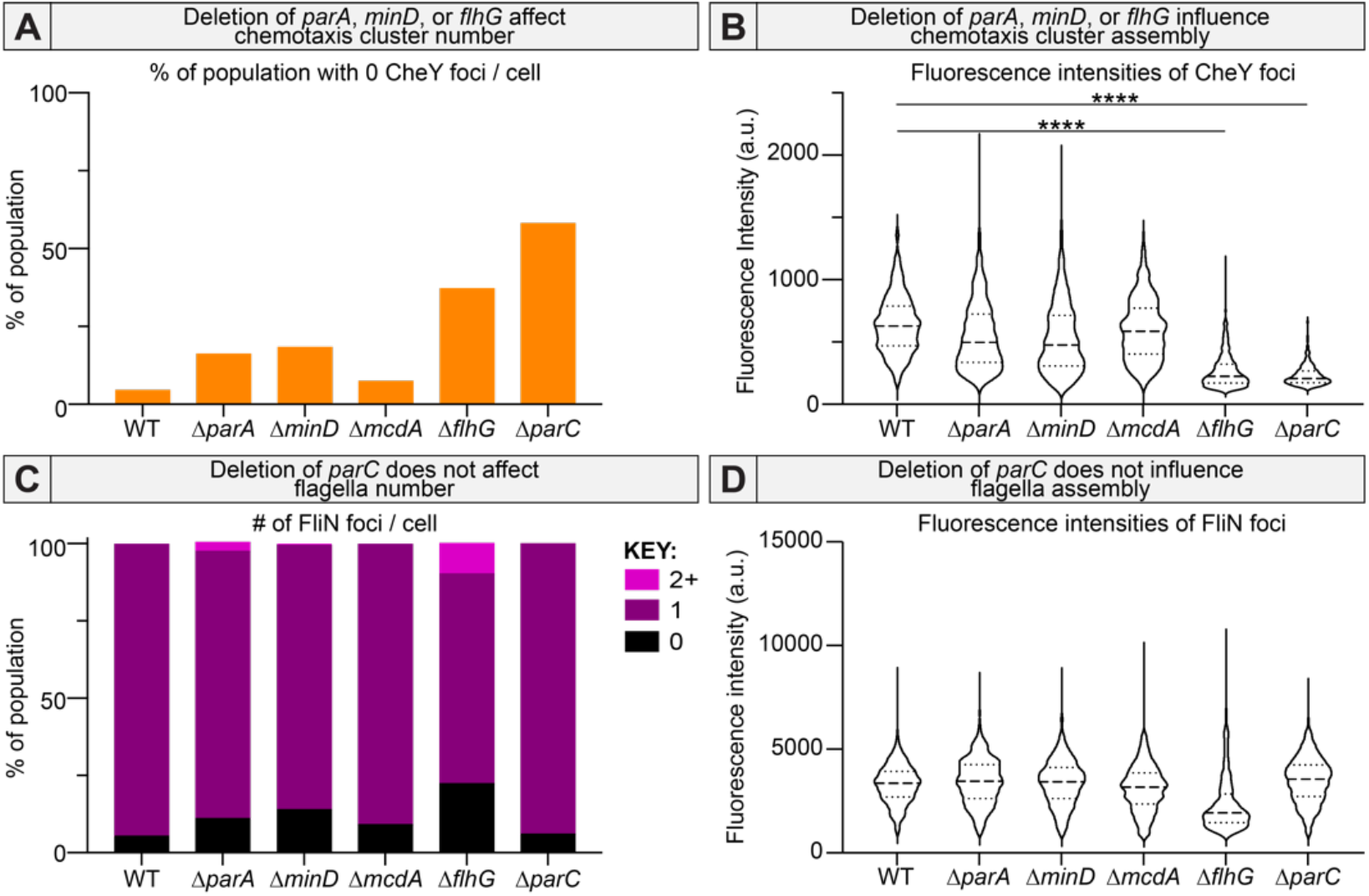
Deletion of ParA, MinD, or FlhG influences chemotaxis cluster. **(A)** number and **(B)** assembly. Only deletion of *flhG* influenced flagella **(C)** number and **(D)** assembly. **** p-value < 0.0001.

It is important to note that while Δ*parA* and Δ*minD* strains exhibited moderate effects on chemotaxis cluster assembly, deletion of *flhG* was more severe **(Figure 6AB)**. Given that chemotaxis clusters communicate with the flagellum to move the bacterium towards favorable conditions, we hypothesize that chemotaxis cluster assembly and organization in *H. neapolitanus* is regulated by flagellum positioning. Interestingly, this effect was not reciprocal, as deletion of *parC* had no effect on flagella foci number **(Figure 6C)** or flagella basal body intensity **(Figure 6D)**. Identifying the molecular players responsible for this crosstalk in the spatial regulation of the flagellum and chemotaxis cluster is a subject of future work. Together, our data demonstrate interdependencies in how A/D ATPases coordinate the positioning of protein-based organelles with each other as well as the processes of DNA segregation and cell division.

### A/D ATPases have unique interfaces that confer cargo-positioning specificity

We have experimentally demonstrated that multiple A/D ATPases coexist and function in the same cell to position multiple disparate cargos. We also showed that A/D-based positioning is cargo specific. A/D ATPases have been shown to form very similar sandwich dimer structures **(Dunham et al., 2009; Leonard et al., 2005; Schumacher et al., 2012; Wu et al., 2011)**, and AlphaFold2 (AF2) predictions suggest this is also the case for all five A/D ATPases of *H. neapolitanus* studied here **(Figure S7)**. The structural similarities suggest there are conserved interfaces unique to each A/D ATPase that provide specificity, linking an A/D ATPase to its particular cargo.

Using AF2 and Rosetta predictions, we determined the A/D ATPase structures from *H. neapolitanus* and identified the putative interaction interfaces that provide specificity to each positioning reaction. There are three interfaces on an A/D ATPase that confers specificity: 1) the dimerization interface, 2) the interface for interaction with its positioning matrix (nucleoid or membrane), and 3) the interface for interaction with its partner protein, which ultimately links the ATPase to its cargo. We identified these three interfaces for all five of the A/D ATPases in *H. neapolitanus* **(Figure 7)** and predict key residues required for these associations **(Table S2)**.

**Figure 7:**
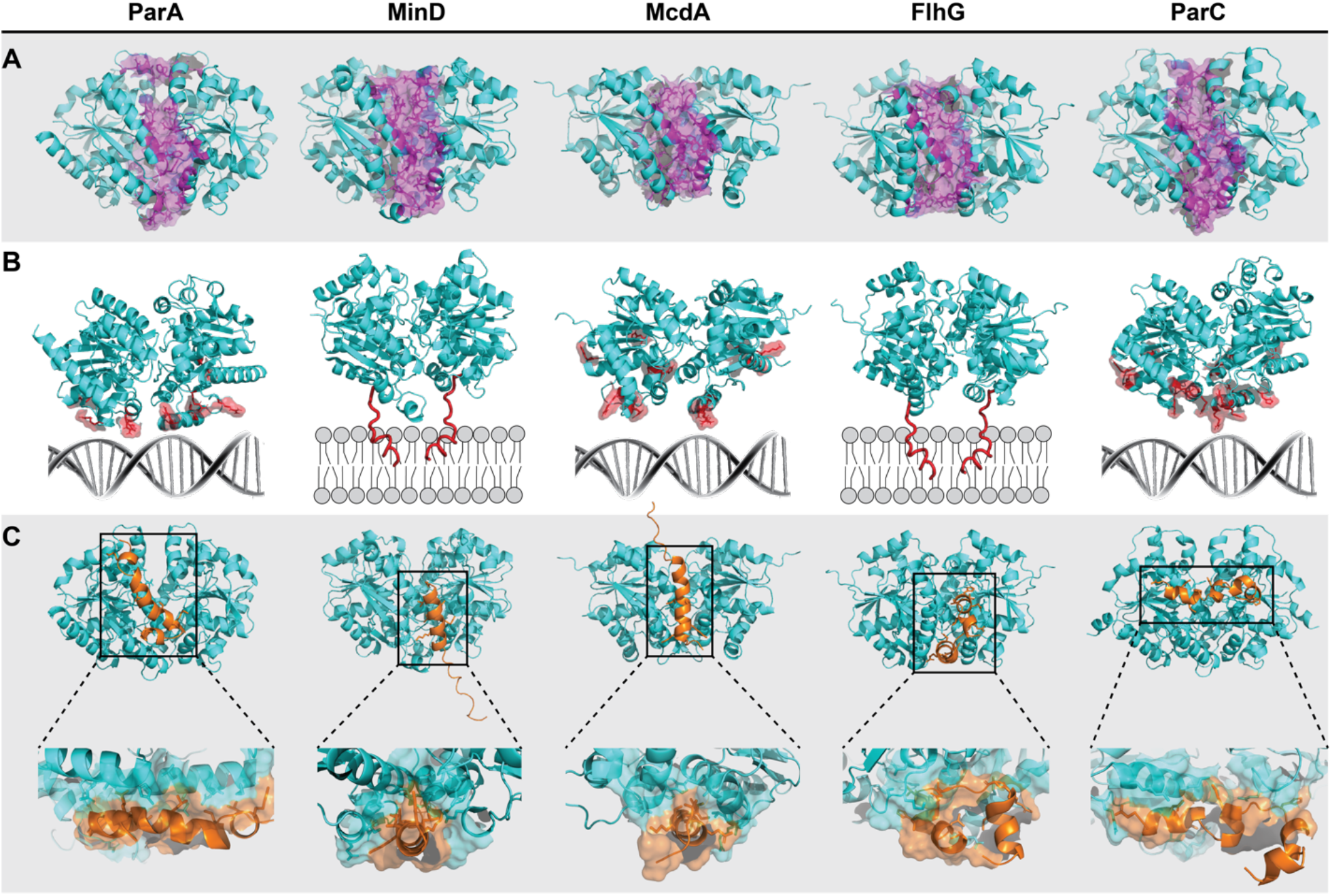
A/D ATPases have unique interfaces that confer cargo-positioning specificity. **(A)** Homodimer structures of the A/D ATPases in *H. neapolitanus* were generated using AlphaFold2 (cyan). Putative residues for homodimer specificity are highlighted magenta. **(B)** Dimers from (A) are oriented over their positioning matrix – nucleoid DNA or membrane. Putative residues critical for binding nucleoid DNA or membrane are highlighted red. **(C)** Dimer structures docked with the N-terminal interacting peptide from putative partner proteins (orange), which confer cargo specificity. Predicted residues critical for this association are space-filled cyan on the ATPase and orange on the partner protein in the zoom.

Specificity at the dimer interface restricts A/D ATPases to homodimerization, and thereby allows each A/D ATPase to function independently in the same cell without cross-interference **(Figure 7A)**. The putative residues required for homodimerization of the five A/D ATPases in *H. neapolitanus* are provided in **Table S2, Tab 1**.

ParA-like ATPases use the nucleoid and MinD-like ATPases use the inner membrane for cargo positioning. ParA, McdA, and ParC have basic residues at their C-termini for non-specific binding to nucleoid DNA, whereas MinD and FlhG have <u>m</u>embrane <u>t</u>argeting <u>s</u>equences (MTSs) for membrane binding **(Figure 7B)**. We identified the predicted residues required for each A/D ATPase to associate with their respective positioning matrix **(Table S2, Tab 2)**.

Cargo specificity for A/D ATPases comes from interacting with a partner protein that either associates with the cargo, or is a physical component of the cargo itself. Partner proteins have a stretch of amino acids enriched in charged residues at the N-terminus that exclusively interacts with its A/D ATPase, while the rest of the partner protein is dedicated to cargo association **(Schumacher et al., 2021)**. We generated docking models of each A/D ATPase with an N-terminal peptide from its partner protein **(Figure 7C)**. The peptides were defined as the first 30 residues of the putative partner protein from the N-terminus. The peptide docking simulations identified several putative residues that are key to system specificity between an A/D ATPase and its partner protein **(Table S2, Tab 3)**.

Finally, we performed *in silico* alanine-scanning mutagenesis across all residues comprising the interacting interface of the N-terminal peptides of partner proteins docked onto their cognate A/D ATPase **(Table S2, Tab 4)**. The resulting ΔΔG values identify the extent to which each residue contributes to the stability of the partner protein interaction with its cognate A/D ATPase. Importantly, our *in silico* alanine-substitution simulations identify the residues experimentally shown to be important for docking the *Escherichia coli* MinE peptide onto the MinD dimer **(Park et al., 2011, 2012; Wu et al., 2011) (Figure S7C)**. Together, the *in silico* data provide a roadmap for strategic mutagenesis and mechanistic probing of the specificity determinants involved in bacterial chromosome segregation, cell division positioning, and protein-based organelle trafficking by A/D ATPases across prokaryotes.

## Discussion

The study of A/D ATPases has focused on a specific cargo of a certain biological process, and largely in model organisms that encode only one or two A/D ATPases. In these studies, two questions are typically posed: How does a specific cargo find its correct position, and how does this position change over time? Here, our focus was on the positioning systems, rather than a specific cargo-type or biological process, and as such, is the first systems biology approach to address how multiple A/D ATPases coordinate the positioning of diverse cargos in the same cell.

### Encoding multiple A/D ATPases is a shared feature across prokaryotes

Over a third of all sequenced bacterial genomes encode multiple A/D ATPases **(Figure 1A)**, with some bacteria encoding more than 10. Interestingly, although most bacteria have the same fundamental cargos, not all use dedicated A/D-based positioning systems for each cargo. For example, many of the cellular cargos we found here to be positioned by A/D ATPases in certain bacteria, like *H. neapolitanus*, are not actively positioned in others, like *E. coli*. What necessitates an A/D ATPase for positioning a certain cellular cargo in one bacterium and not in another remains an open question. There does, however, seem to be a limit to the number of A/D ATPases that a bacterium can encode.

A/D ATPases are also encoded in archaeal genomes **(Leipe et al., 2002)**, but little is known about their roles in subcellular organization. A recent study showed that archaeal species across several phyla, Euryarchaeota in particular, encode multiple A/D ATPases **(Nußbaum et al., 2020)**. Several of these species contained more than a dozen, including *H. volcanii* with 13 A/D ATPases, four of which are MinD-homologs. Strikingly, all four MinD homologs were not required for cell division, but one (MinD4) stimulated the formation of chemotaxis arrays and the archaella, which is the functional equivalent of the bacterial flagellum. This study stresses the importance of experimentally linking A/D ATPases to their cellular cargos as we show here.

### 5 A/D ATPases, 5 Cargos, 1 Cell

The ParA/MinD (A/D) family of ATPases spatiotemporally regulates a growing list of diverse mesoscale complexes critical to fundamental processes in prokaryotes, including cell growth and division, DNA segregation, motility, conjugation, and pathogenesis **(Lutkenhaus, 2012; Vecchiarelli et al., 2012)**. Therefore, understanding how A/D ATPases coordinate the positioning of cellular cargos at the correct location at the correct time is key to understanding bacterial cell function. Using *H. neapolitanus* as a non-pathogenic model, our findings strongly support the idea that each ATPase is dedicated to the positioning of a specific cellular cargo. Our model also unveiled the dependency of protein-based organelle trafficking on the faithful replication and segregation of the chromosome as well as positioning cell division at mid-cell. For example, while the direct consequence of deleting ParA was chromosome missegregation, the resulting asymmetric chromosome inheritance also led to indirect defects in the positioning of carboxysomes and chemotaxis clusters. We also identified a previously uncharacterized epistatic relationship in flagella positioning by FlhG and its downstream influence on the spatial regulation of the chemotaxis cluster by ParC. It is well known that chemotaxis clusters direct cell motility by controlling the direction of flagellar rotation. However, to our knowledge, this is the first study showing crosstalk in the spatial regulation of flagella and chemotaxis clusters. Future studies will determine the molecular players involved in the crosstalk between A/D-based positioning of these two cellular cargos both involved in cell motility.

Our bioinformatics also identified a 6^th^ A/D ATPase in the *H. neapolitanus* genome. Gene *Hn1669* is an A/D ATPase that shows homology to VirC1 and is located near *trb* genes, which encode conjugation machinery components **(Figure 1B)**. A single study has shown that the VirC1 ATPase, encoded on the Ti plasmid of *Agrobacterium tumefaciens*, is involved in recruiting the conjugative Ti plasmid to the Type IV secretion machine at the cell poles **(Atmakuri et al., 2007)**. Intriguingly, in *H. neapolitanus*, the VirC1 homolog is encoded on an Integrative Conjugative Element (ICE) and the downstream gene from this A/D ATPase encodes a ParB homolog. ICEs are major drivers of bacterial evolution and the spread of antibiotic resistance genes (Norman et al., 2009). We have yet to image conjugation in *H. neapolitanus*, but we suspect this ParB homolog binds and demarcates the ICE locus during conjugation. It is attractive to speculate that the VirC1-homolog and its downstream ParB homolog are involved in transporting and positioning the ICE locus to the Type IV secretion machine at the cell pole for conjugation.

### Partner proteins and specificity determinants linking an A/D ATPase to its cargo

While A/D ATPases share sequence, structural, and biochemical commonalities, the partner proteins linking these ATPases to their cargos are extremely diverse. Due to this diversity, the partner protein has not always been identified, and as a result, many A/D ATPases are called ‘orphans’ **(Lutkenhaus, 2012)**. The extreme diversity is largely due to the partner proteins providing the specificity determinants linking an A/D ATPase to its cognate cargo. Despite their extreme diversity, data across the field supports the idea that partner proteins interact with and stimulate their A/D ATPases via a shared mechanism. Partner proteins involved in plasmid partition and chromosome segregation, as well as those required for positioning BMCs, flagella, chemotaxis clusters, and the divisome have all been shown, or suggested, to interact with their A/D ATPase via a positively charged and disordered N-terminus **(Schumacher et al., 2021)**. Our *in silico* analysis of A/D ATPase dimers docked with N-terminal peptides of their partner proteins demonstrates how specific cargos are assigned, and how these related positioning systems coexist and function in the same cell without cross-interference.

### Future Directions

Going forward, we aim to use *H. neapolitanus* as a model to define the general mode of transport shared among the entire A/D ATPase family and to determine how positioning reactions are altered for disparate cargos. These findings are significant because A/D ATPases spatially organize essentially all aspects of bacterial cell function. An additional future direction is to experimentally verify the specificity determinants we identified here for each partner protein and cargo, and leverage this knowledge in the rational design of positioning systems in bacteria. These contributions are expected to be significant because <u>m</u>inimal self-organizing systems are vital tools for synthetic biology **(Schwille and Diez, 2009)**. We aim to design <u>M</u>inimal <u>A</u>utonomous Positioning <u>S</u>ystems (MAPS) consisting of a positioning ATPase and their partner-protein N-terminal peptide as a “luggage tag” to be used as spatial regulators for natural-and synthetic-cargos in heterologous bacteria.

## STAR Methods

### tBLASTn analysis

tBLASTn analysis was done using a ParA/MinD consensus sequence, as a query against RefSeq Representative genomes database with max target sequences as 5000 and E value threshold at 0.0001. Sequences were filtered for those that shared sequence homology and had one of the identified putative cargo genes, confirmed using webFlaGs (https://pubmed.ncbi.nlm.nih.gov/32956448/). The consensus query was generated using COBOLT (*“https://pubmed.ncbi.nlm.nih.gov/17332019/”*).

### FlaG analysis

A few representative genomes were selected to display gene neighborhood conservation. Identification of replication origins (OriC’s) were performed using Ori-Finder (https://bmcbioinformatics.biomedcentral.com/articles/10.1186/1471-2105-9-79). FlaGs analysis figure was generated using Gene Graphics (https://katlabs.cc/genegraphics/).

### Multiple sequence alignment

Sequences were aligned using Clustal Omega. The resulting tree was imported into iTOL to generate an unrooted tree. (Accession numbers: Hn2335 – ACX97145.1; Hn1364 – ACX96198.1; Hn0912 – ACX95755.1; Hn0716 – ACX95565.1; Hn0722 – ACX95571.1; Hn1669 – ACX96495.1; Hn0255 – ACX95118.1)

### Media and growth conditions

All mutants described in this study were constructed using WT *Halothiobacillus neapolitanus* (Parker) Kelly and Wood (ATCC® 23641™) purchased from ATCC. Cultures were grown in ATCC® Medium 290: S-6 medium for Thiobacilli (Hutchinson et al., 1965) and incubated at 30°C, while shaken at 130 RPM in air supplemented with 5% CO_2_. Strains were preserved frozen at −80°C in 10% DMSO.

### Construct designs and cloning

All constructs were generated using Gibson Assembly and verified by sequencing. Fragments for assembly were synthesized by PCR or ordered as a gBlock (IDT). Constructs contained flanking DNA that ranged from 750 to 1100 bp in length upstream and downstream of the targeted insertion site to promote homologous recombination into target genomic loci. Cloning of plasmids was performed in chemically competent *E. coli* Top10 or Stellar cells (Takara Bio).

### Making competent cells in H. neapolitanus C2

Competent cells of *H. neapolitanus* were generated as previously reported. In short, 1 L of culture was grown to an OD of 0.1-0.15. Cultures were harvested by centrifugation at 5,000xg for 20 minutes at 4°C. Pellets were resuspended and washed twice with 0.5 volumes of ice-cold nanopore water. All wash centrifugation steps were performed at 3,000xg for 30 minutes at 4°C. The resulting pellet after washing was resuspended in 1×10^−3^ volumes of ice-cold nanopore water. These competent cells were used immediately or frozen at -80°C for future use. Frozen competent cells were thawed at 4°C before use.

### Transformation in H. neapolitanus C2

50-100 µL of competent cells were mixed with 5 µL plasmid DNA (1-5 µg) and incubated on ice for 5 minutes. This mixture was then transferred to a tube containing 5 mL ice-cold S6 medium without antibiotics and incubated on ice for 5 minutes. Transformations were recovered for 16-36 hours, while shaken at 130 RPM, at 30°C, in air supplemented with 5% CO_2_. Clones were selected by plating on selective medium with antibiotics. Colonies were restreaked. Restreaked colonies were verified for mutation by PCR.

### Native fluorescent fusions

For the native fluorescent fusion of ParB-mNG, mNG-FliN, and CheY-mNG, the sequence encoding the fluorescent protein mNeonGreen (mNG) was attached to the 3’ or 5’ region of the native coding sequences, separated by a GSGSGS linker. For the native fluorescent fusion of Cbbs-mTQ, the sequence encoding the fluorescent protein mTurquoise (mTQ) was attached to the 3’ region of the native coding sequence, separated by a GSGSGS linker. A kanamycin resistance cassette was inserted before the gene for N-terminal tags or after the gene for C-terminal tags. When necessary, the promoter was duplicated. The mutant was selected by plating on S6 agar plates supplemented with 50 μg/mL of kanamycin. All fusions were verified by PCR.

### Deletion mutants

For deletions of *Hn2335, Hn1364, Hn0912*, and *Hn0722*, the genes were replaced with a spectinomycin resistance cassette, followed by a duplicated promoter for the downstream gene. Deletion of *Hn0716* was obtained by codon-optimizing the downstream gene and inserting the spectinomycin resistance cassette after this codon-optimized gene. Mutants were selected by plating on S6 agar plates supplemented with 50 μg/mL of spectinomycin. All mutations were verified by PCR.

### Microscopy

All live-cell microscopy was performed using exponentially growing cells. 3-5 µL of cells were dropped onto a piece of 2% UltraPure agarose + S6 pad and imaged on a Mantek dish. All fluorescence and phase contrast imaging were performed using a Nikon Ti2-E motorized inverted microscope controlled by NIS Elements software with a SOLA 365 LED light source, a 100X Objective lens (Oil CFI Plan Apochromat DM Lambda Series for Phase Contrast), and a Photometrics Prime 95B Back-illuminated sCMOS camera or a Hamamatsu Orca Flash 4.0 LT + sCMOS camera. ParB-mNG, mNG-FliN, and CheY-mNG were imaged using a “GFP” filter set (C-FL GFP, Hard Coat, High Signal-to-Noise, Zero Shift, Excitation: 470/40 nm [450-490 nm], Emission: 525/50nm [500-550nm], Dichroic Mirror: 495 nm). CbbS-mTQ labeled carboxysomes were imaged using a “CFP” filter set (C-FL CFP, Hard Coat, High Signal-to-Noise, Zero Shift, Excitation: 436/20 nm [426-446 nm], Emission: 480/40 nm [460-500 nm], Dichroic Mirror: 455 nm). DAPI fluorescence was imaged using a standard “DAPI” filter set (C-FL DAPI, Hard Coat, High Signal-to-Noise, Zero Shift, Excitation: 350/50 nm [325-375 nm], Emission: 460/50 nm [435-485 nm], Dichroic Mirror: 400 nm). Alexa Fluor 594 C5 maleimide-conjugated flagella were imaged using a “TexasRed” filter set (C-FL Texas Red, Hard Coat, High Signal-to-Noise, Zero Shift, Excitation: 560/40 nm [540-580 nm], Emission: 630/75 nm [593-668 nm], Dichroic Mirror: 585 nm).

### Long-term microscopy

For multigenerational time-lapse microscopy, 1.5% UltraPure agarose + S6 pads were cast in 35-mm glass-bottom dishes. Dishes were preincubated at 30°C in 5% CO_2_ for at least 24 hr. 4 µl of exponentially growing cells were spotted onto the agar pad. Temperature, humidity, and CO_2_ concentrations were controlled with a Tokai Hit Incubation System. NIS Elements software was used for image acquisition. Cells were preincubated in the stage top for at least 30 minutes before image acquisition. Videos were taken at one frame per 2.5-5 minutes for a duration of 12-24 hours.

### Image analysis

Several fields of view were captured for each mutant. All fluorescence channels were subjected to background subtraction on Fiji with a rolling ball radius of 50 μm. Background-subtracted fluorescence images were merged with phase contrast images to create composites used for image analysis. Image analysis including cell identification, quantification of cell length, foci localization, foci number, foci fluorescence intensity, and identification of constriction sites were performed using Fiji plugin MicrobeJ 5.13I (Schindelin et al., 2012; Ducret et al., 2016). Cell perimeter detection and segmentation were done using the rod-shaped descriptor with default threshold settings at a tolerance of 56. Maxima detection parameters were individually set for each cargo. For ParB-mNG (cargo = chromosome) foci detection, tolerance and z-score were both set to 100. For CheY-mNG (cargo = chemotaxis) foci detection, tolerance was set to 150 and z-score was set to 100. For mNG-FliN (cargo = flagella) foci detection, tolerance was set to 710 and z-score was set to 69. For CbbS-mTQ (cargo = carboxysome) foci detection, point detection was used instead of foci detection, and tolerance was set to 1010. Results were manually verified using the experiment editor, and non-segmented cells were cut using a particle cutter. Associations, shape descriptors, profiles, and localization were recorded for each strain. Localization graphs were automatically generated through MicrobeJ. Fluorescence intensity graphs and foci number count graphs were made in GraphPad Prism (GraphPad Software, San Diego, CA, www.graphpad.com).

### Nucleoid staining for live imaging

Cells were harvested by centrifugation at 5,000 g for 5 minutes. Following centrifugation, the cells were washed in PBS, pH 7.4. The resulting cell pellet was resuspended in 100 μL PBS and stained with SytoxBlue at a final concentration of 500 nM. The samples were incubated in the dark, at room temperature for 5-10 minutes. Stained cells were directly loaded onto an S6 agar pad that had been infused with 500 nM SytoxBlue.

### Carboxysome count

Eleven frames were captured over the course of 2 minutes. Carboxysomes were counted in each frame. The highest number counted per cell was used for carboxysome count.

### Identifying constriction sites

For each mutant strain, ∼200 dividing cells were analyzed for the location of their constriction sites relative to mid-cell.

### Motility assay

Motility assays were run in S6 in 0.4% agar. Cells were grown on a plate of S6 medium. Individual colonies were inoculated into tubes or plates of motility media and incubated at 30°C in air supplemented with 5% CO_2_. Tubes and plates were checked daily for motility for 2 weeks.

### Cysteine-labelling of flagellin

*H. neapolitanus* flagellin was identified using a BLAST homology search with Hag from *Bacillus subtilis* as the query. Only a single flagellin gene was found. Several threonine and serine residues were considered for cysteine mutagenesis. Residues were mutated using the Q5 site-directed mutagenesis kit. Clones were verified for cysteine mutation by sequencing.

### Flagella stain

Alexa Fluor 594 C5 maleimide dye was resuspended in DMSO to the working concentration of 10 mM. *H. neapolitanus* cultures were grown to an OD of 0.1-0.2. The cultures were adjusted to a pH of 7.0 using PBS, pH 11.7. Cells at an adjusted pH of 7.0 were then stained with Alexa Fluor 594 C5 maleimide dye at a final dye concentration of 100 µM. Staining cultures were incubated overnight at 4°C. The stained cells were washed 4+ times in PBS, pH 7.4. All centrifugation steps were performed at 5,000xg for 3 minutes.

### Statistical analysis

For all population analyses, we used a non-parametric Wilcoxon test using GraphPad Prism.

### Movie editing

Movies were cropped using Fiji. Movies were stabilized using the multi-channel hyperstack alignment plug-in called HyperStackReg. Time stamp and scale bar annotations were added using Fiji. Arrows and pauses were added using Adobe Premier Pro.

### Structure prediction and peptide docking using AlphaFold2 and Rosetta

For MinD and McdA, we generated the N-terminus peptide-ATPase docking models using the CollabFold implementation of AlphaFold2 **(Jumper et al., 2021; Mirdita et al., 2022)**. The N-terminal peptides were defined as the first 30 residues of the putative partner protein from the N-terminus. Multiple sequence alignments were constructed using MMseqs2. For each peptide-ATPase pair, we generated five structures with the default CollabFold/AlphaFold2 hyperparameters saved for the number of recycles for each model being increased to 12. As default to Alphafold2, the structures were energetically minimized with AMBER using the Amber99sb force field. We selected docked peptide models based on the pLDDT scores of the binding interface residues and similarity to previously resolved ParA-like ATPase/partner-protein crystal structures.

For ParA, FlhG, and ParC we generated docked peptide models using Rosetta’s FlexPepDock protocol **(Raveh et al., 2011)**. First, we equilibrated the ATPase homodimer structures generated from AlphaFold2 to the ref2015_cart_cst Rosetta force field with the FastRelax full-atom refinement protocol with cartesian coordinate space minimization using the lbfgs_armijo_nonmonotone minimizer. To preserve the position of the backbone atoms predicted by AlphaFold2, a backbone atom coordinate constraint was added. For this initial step, we generated 20 trajectories with the lowest scoring structure being used for the peptide docking step. We used the score3 with the docking_cen.wts_patch and the REF2015 force fields for the low-resolution and high-resolution docking steps of the FlexPepDock protocol, respectively. For each putative partner protein/ATPase pair, we simulated 50,000 docking trajectories. The top two thousand trajectories defined by the lowest energetics were then clustered using Calibur (Li and Ng, 2010). We then chose the final models based on cluster information, energetics, and mechanistic plausibility. From the final docked models chosen from the Rosetta and Alphafold2 simulations, we sought to identify key binding residues through an *in-silico* alanine mutation scan and ΔΔG calculations. We iteratively mutated all interface residues of the docked peptide and calculated the ΔΔG using the FlexDDG protocol in Rosetta with the talaris2014 forcefield **(Barlow et al., 2018)**. In FlexDDG, for each mutation, the backbone and side chain conformations were sampled 35,000 times using Rosetta’s Monte Carlo backrub method. At every 2,500 sample interval the ΔΔG of mutation was calculated. The final reported ΔΔG is the average of 35 such trajectories.

(Accession numbers: ParA partner protein (ParB) - **ACX97144.1;** MinD partner protein (MinE) - **ACX96199.1**; McdA partner protein (McdB) - **ACX95754.1;** FlhG partner protein (FliA) - **ACX95566.1;** ParC partner protein (CheW) - **ACX95572.1)**. The first 30 amino acids from the N-terminus of each partner protein were docked onto the dimer structures *in silico* using AlphaFold2 (MinD, McdA) or Rosetta (ParA, FlhG, ParC). Binding interface was defined by those residues that shared a minimum of 1 Angstrom^2^ of surface area.

## Data Availability

All data generated or analyzed during this study are included in this published article [and its supplementary information files].

## Acknowledgements

We thank Joshua S. MacCready, Christopher A. Azaldegui, Joseph L. Basalla, Ajai J. Pulianmackal, and Claudia Mak for thoughtful discussions. We thank Holly Turula, Maria Ghalmi, and Miles Mckenna for their technical assistance. This work was supported by the National Science Foundation to A.G.V (Award No. 1817478), NSF GRFP to L.T.P. (DGE 1841052), and from research initiation funds provided by the MCDB Department to A.G.V.

## Author Contributions

Conceptualization, L.T.P. and A.G.V.; Methodology, L.T.P., M.J.L., K.R.; Formal Analysis, L.T.P. M.J.L., K.R., S.Y., M.J.O. and A.G.V.; Investigation, L.T.P., J.Z., and M.K.T.; Resources, A.G.V.; Writing – Original Draft, L.T.P. and A.G.V.; Visualization, L.T.P., M.J.L., K.R., and A.G.V.; Supervision, L.T.P., M.J.O., and A.G.V.; Funding Acquisition, L.T.P. and A.G.V.

---

**Figure S1:**
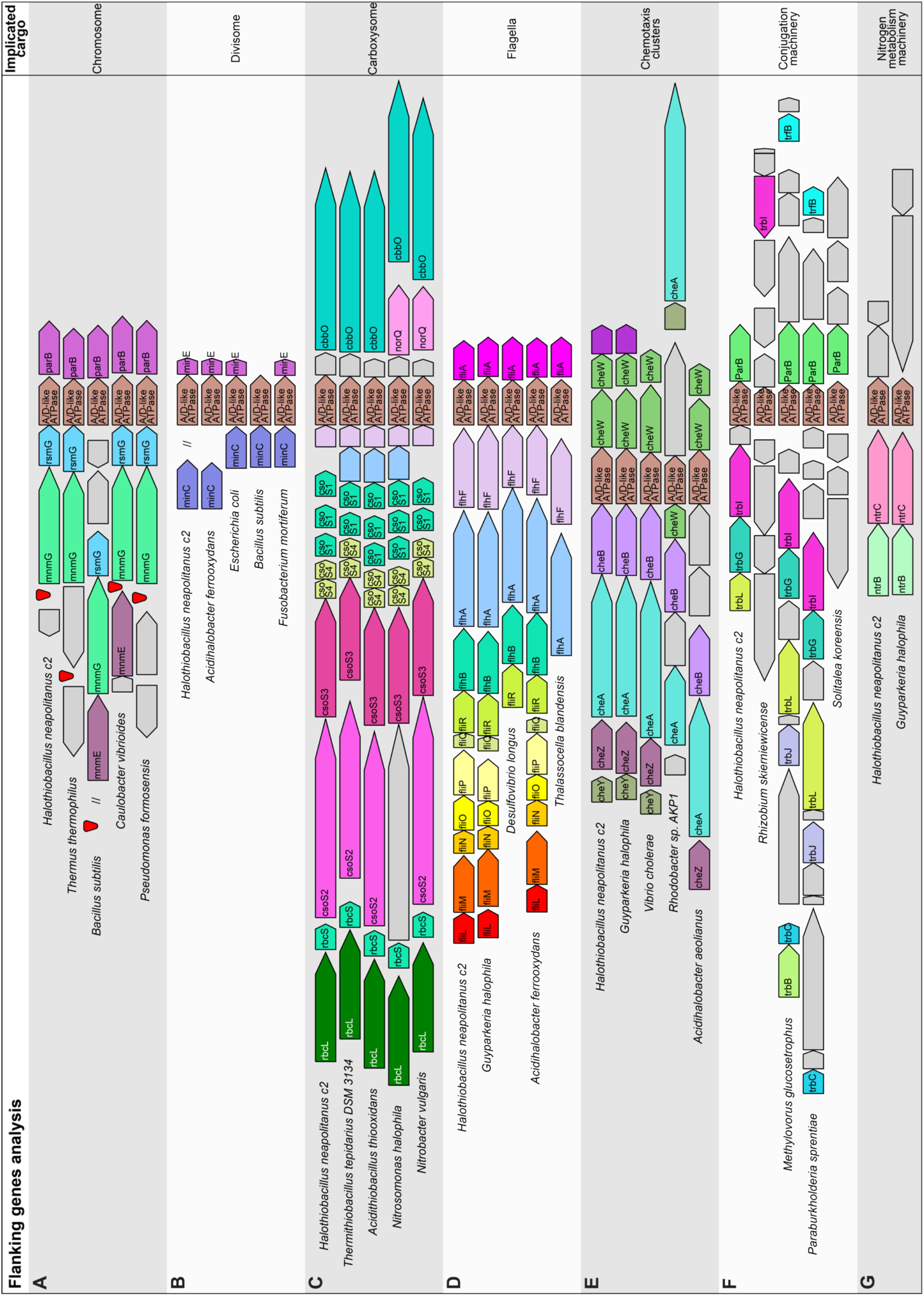
<u>Fla</u>nking <u>G</u>enes (FlaGs) analysis shows conservation among A/D ATPase gene neighborhoods. **(A)** The A/D ATPase gene involved in chromosome segregation (*parA*) is typically found a few genes downstream of the origin of replication (red cone) and upstream of a *parB* gene. **(B)** The A/D ATPase gene involved in divisome positioning (*minD*) is typically between genes encoding MinC and MinE. MinC in *H. neapolitanus* was found elsewhere in the genome. **(C)** The A/D ATPase gene involved in carboxysome distribution (*mcdA*) is typically found near carboxysome shell components. **(D)** The A/D ATPase gene involved in flagella positioning (*flhG*) is typically found near core components of the flagellar apparatus. **(E)** The A/D ATPase gene involved in chemotaxis cluster positioning (*parC*) is typically found near genes required for chemotaxis cluster formation and regulation. **(F)** The gene neighborhoods of conjugation operons are not well conserved. However, the A/D ATPase gene involved in spatially organizing conjugation (*virC1*) is typically found upstream of a ParB-like protein. The conjugation operons in *Methylovorus glucosetrophus* and *Paraburkholderia sprentiae* were found on plasmids pMsip01 and pI3WSM5005, respectively. **(G)** Gene neighborhood of A/D ATPase genes associated with nitrogen metabolism are not well conserved. Only one other organism had similar neighboring genes as *H. neapolitanus*.

**Figure S2:**
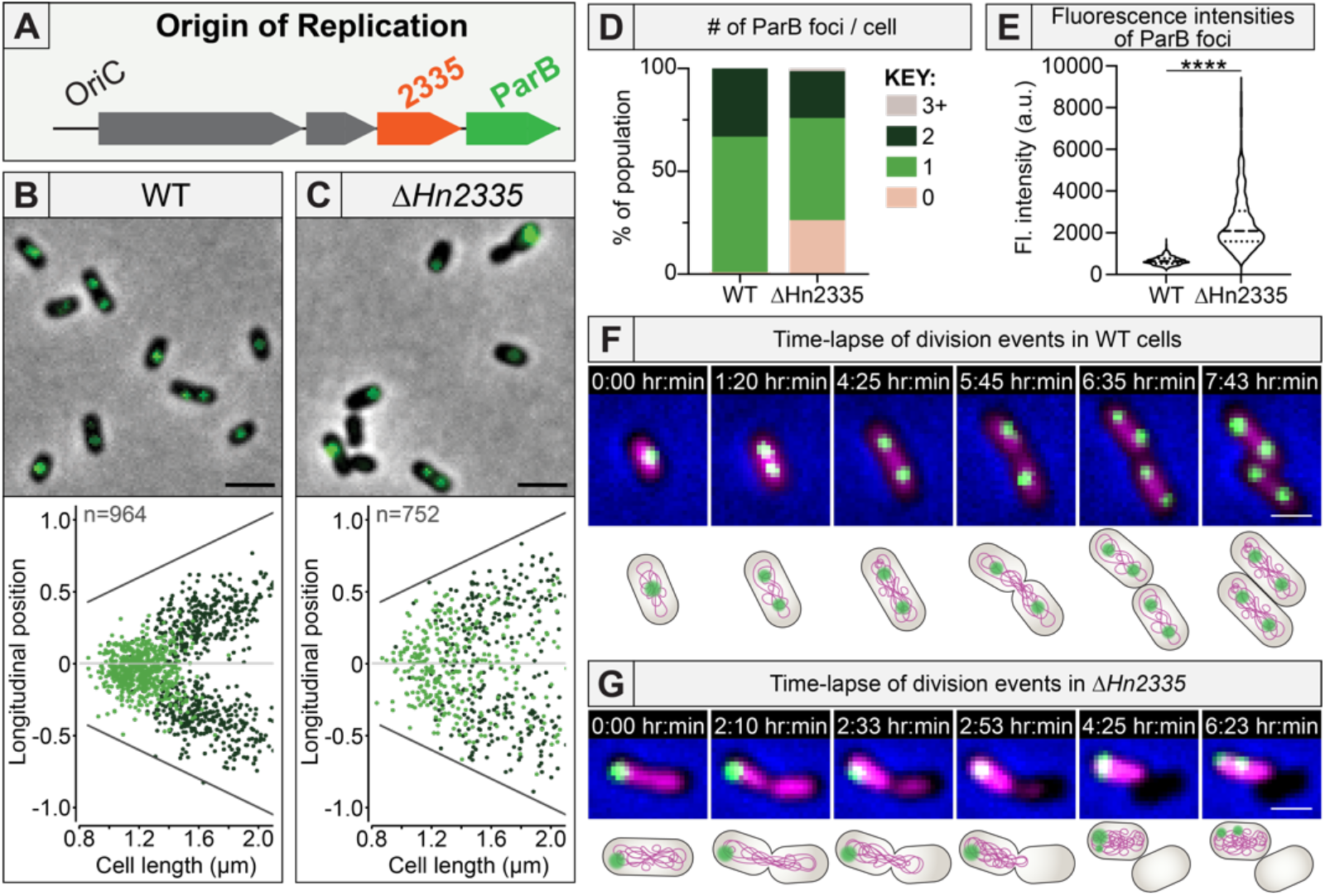
*Hn2335* is required for chromosome segregation in *H. neapolitanus*. **(A)** *Hn2335* is found near the origin of replication (*OriC*) and has a ParB-homolog encoded immediately downstream. The genomic location of *Hn2335* suggests it encodes for the chromosome segregation ParA ATPase. **(B)** The chromosome origin of replication was tagged by labelling the ParB homolog with <u>m</u>Neon<u>G</u>reen to form ParB-mNG. Population analysis of foci localization: Cells were analyzed and quantified using MicrobeJ. On the x-axis, cells are organized according to increasing cell length. The y-axis represents the distance from mid-cell (μm). The foci on the graphs represent where the ParB foci are found along the length of the cell. Light green: 1 focus/cell; dark green: 2 foci/cell. Short WT cells had a single ParB focus at mid-cell, whereas longer cells had two foci at the quarter positions. Scale bar: 2 μm **(C)** Δ*Hn2335* mutant cells displayed random positioning of ParB foci regardless of cell length. Scale bar: 2 μm **(D)** WT cells had 1-2 foci. 25% of Δ*Hn2335* cells had no foci. **(E)** Δ*Hn2335* cells had much brighter ParB foci compared to WT. Wilcoxon test p-value < 0.0001 **(F)** Newborn WT cells have a single ParB focus at mid-cell. Foci are then faithfully segregated to the quarter positions prior to division. Green: ParB foci; Magenta: SytoxBlue stain. Scale bar: 1 μm **(G)** In Δ*Hn2335*, faithful foci positioning and segregation is lost. Because both ParB foci remain on the left-hand side of the dividing cell, the cell on the right becomes anucleate following cell division. Green: ParB foci; Magenta: SytoxBlue nucleoid stain. Scale bar: 1 μm

**Figure S3:**
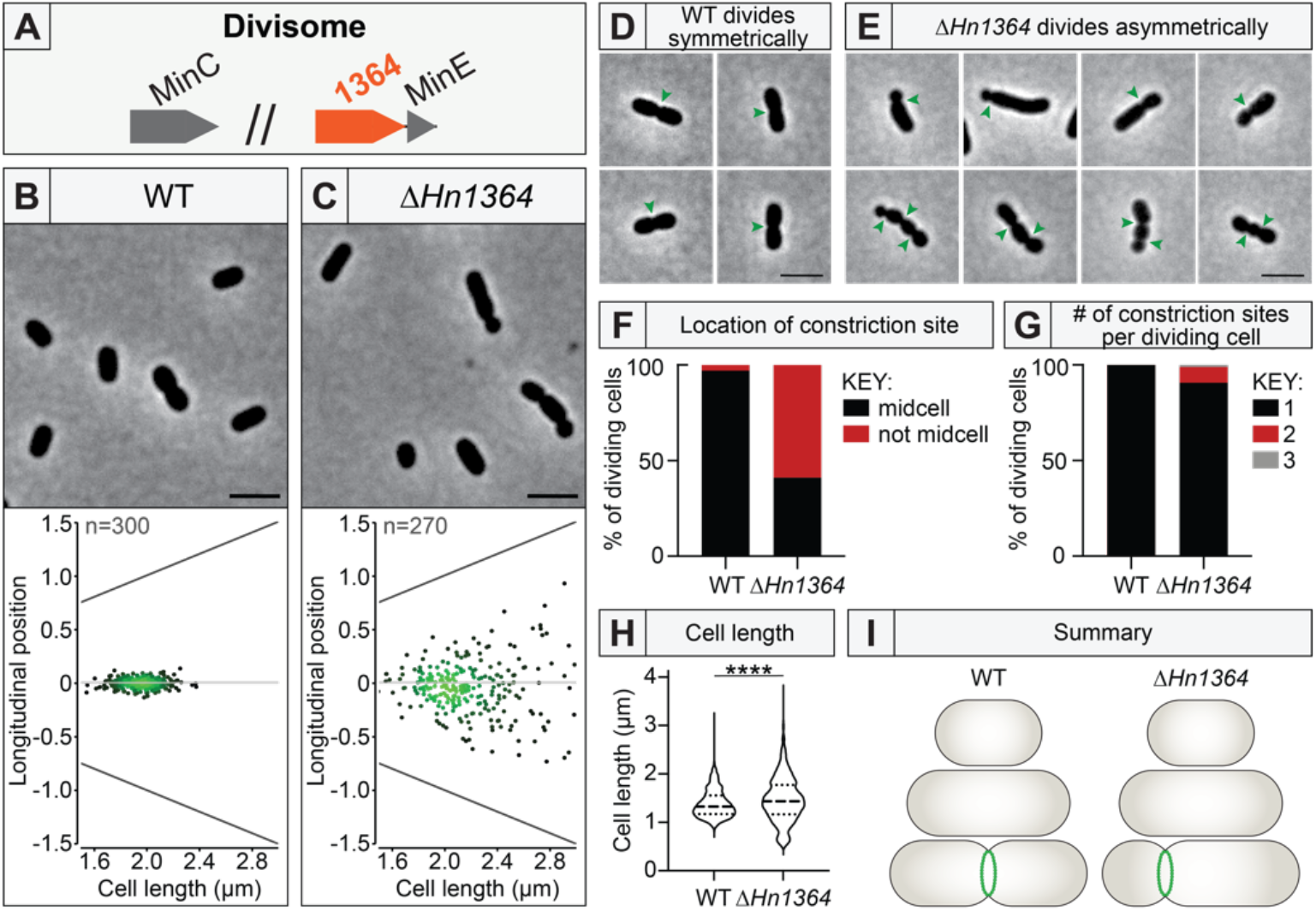
*Hn1364* is required for cell division positioning. **(A)** *Hn1364* is found directly upstream of *minE. minC* is also present elsewhere in the genome. Therefore, the MinCDE system is present and likely involved in divisome positioning in *H. neapolitanus*. **(B, C)** Divisome positioning was determined by the location of constriction sites. Cells were analyzed and quantified using MicrobeJ. For each mutant, ∼300 dividing cells were analyzed for the location of their constriction sites relative to mid-cell. On the x-axis, cells are organized according to increasing cell length. The y-axis represents the distance from mid-cell (μm). Each dot on the graph represents one identified constriction site. In this density plot, light green represents high density and dark green represents low density. In WT cells, constriction sites were found close to mid-cell. In Δ*Hn1364*, constriction sites could be found throughout the length of the cell. Scale bar: 2 μm **(D)** WT cells had constriction sites at mid-cell (green arrows). Scale bar: 2 μm **(E)** Δ*Hn1364* cells were more likely to divide asymmetrically at non-mid-cell locations (green arrows). Multiple division sites could also be found simultaneously on the same cell. Scale bar: 2 μm **(F)** 97% of WT cells divided at mid-cell. Only 41% of Δ*Hn1364* cells divided at mid-cell. Constriction sites were considered “mid-cell” when they were found within 5% of the cell center along the long axis. **(G)** WT cells only had one division site per cell at any given time. In Δ*Hn1364*, 9% of dividing cells had multiple division sites per cell. **(H)** Mutant cells displayed greater variability in cell size. Wilcoxon test p-value < 0.0001 **(I)** *Hn1364* is critical for positioning the divisome at mid-cell in *H. neapolitanus*.

**Figure S4:**
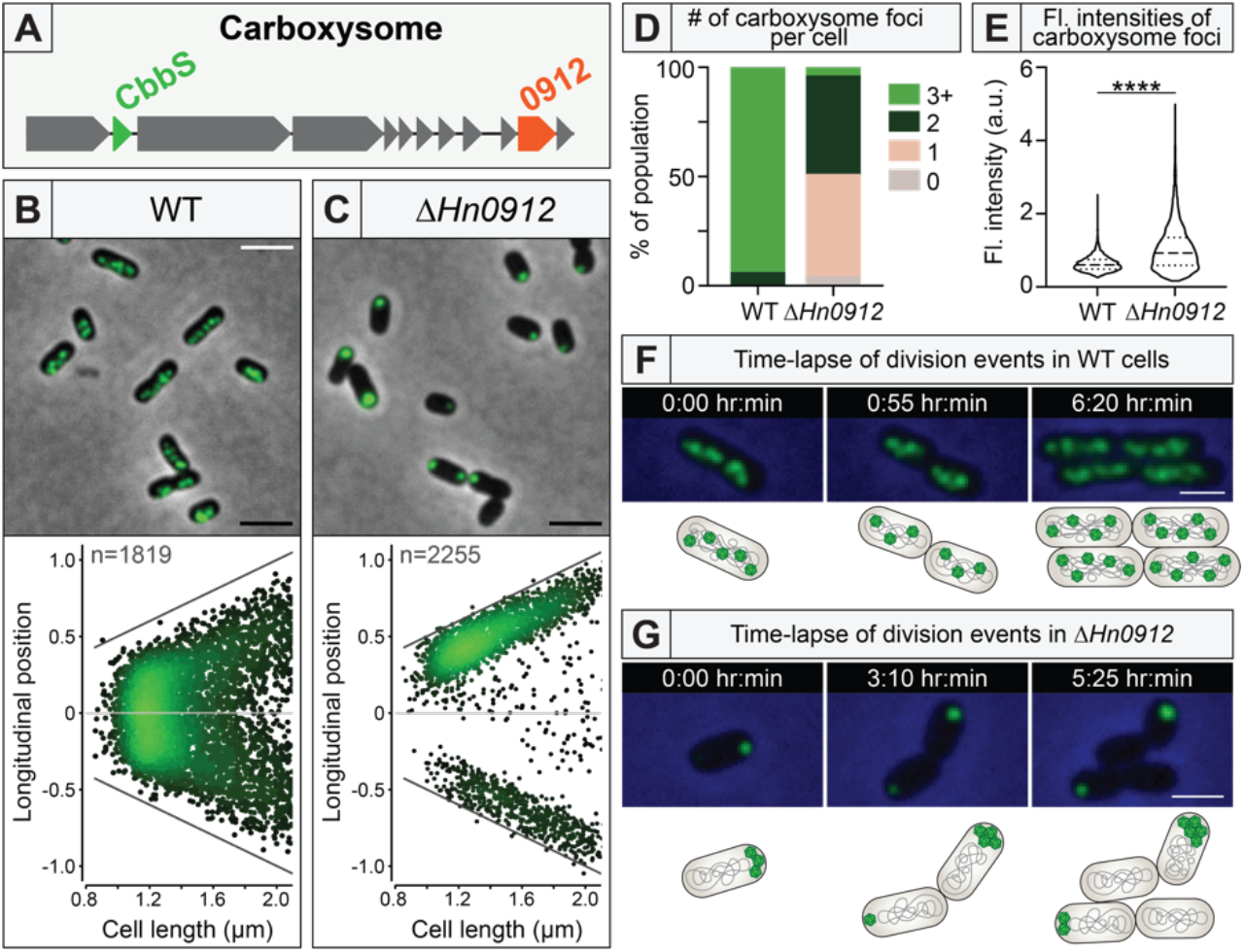
Carboxysome positioning is determined by McdA, the ParA/MinD-like ATPase encoded in the carboxysome operon. **(A)** *Hn0912* encodes McdA and is found near genes encoding carboxysome shell proteins and RuBisCO. **(B)** Carboxysomes were visualized by labelling the small subunit of the Rubisco enzyme (*cbbS*) with mTurquoise2 to form CbbS-mTQ. Population analysis of foci localization: Cells were analyzed and quantified using MicrobeJ. On the x-axis, cells are organized according to increasing cell length. The y-axis represents the distance from mid-cell (μm). The foci on the graphs represent where the foci are found along the length of the cell. In WT cells, carboxysomes are distributed across the cell length. Scale bar: 2 μm **(C)** In Δ*mcdA*, carboxysomes formed a large polar focus at one or both poles. The pole of the cell that is closest to a focus is oriented to the top. Foci in the bottom half of the graph indicate a second focus. Scale bar: 2 μm **(D)** Carboxysome foci count varied dramatically between WT and mutant. In WT, 95% of cells had three or more foci, compared to only 5% of mutant cells. Instead, the vast majority of mutant cells had one or two carboxysome foci at the cell poles. Additionally, ∼ 5% of the mutant population had no foci, which was never observed in WT cells. **(E)** Fluorescence intensity analysis of the foci revealed that, although the mutant population had fewer foci, the foci were significantly brighter. Wilcoxon test p-value < 0.0001 **(F)** In WT, carboxysomes are dynamically positioned along the cell length throughout the cell cycle and across multiple generations. Scale bar: 2 μm **(G)** In Δ*mcdA*, carboxysome aggregates were stagnant throughout the cell cycle and across multiple generations. Scale bar: 2 μm

**Figure S5:**
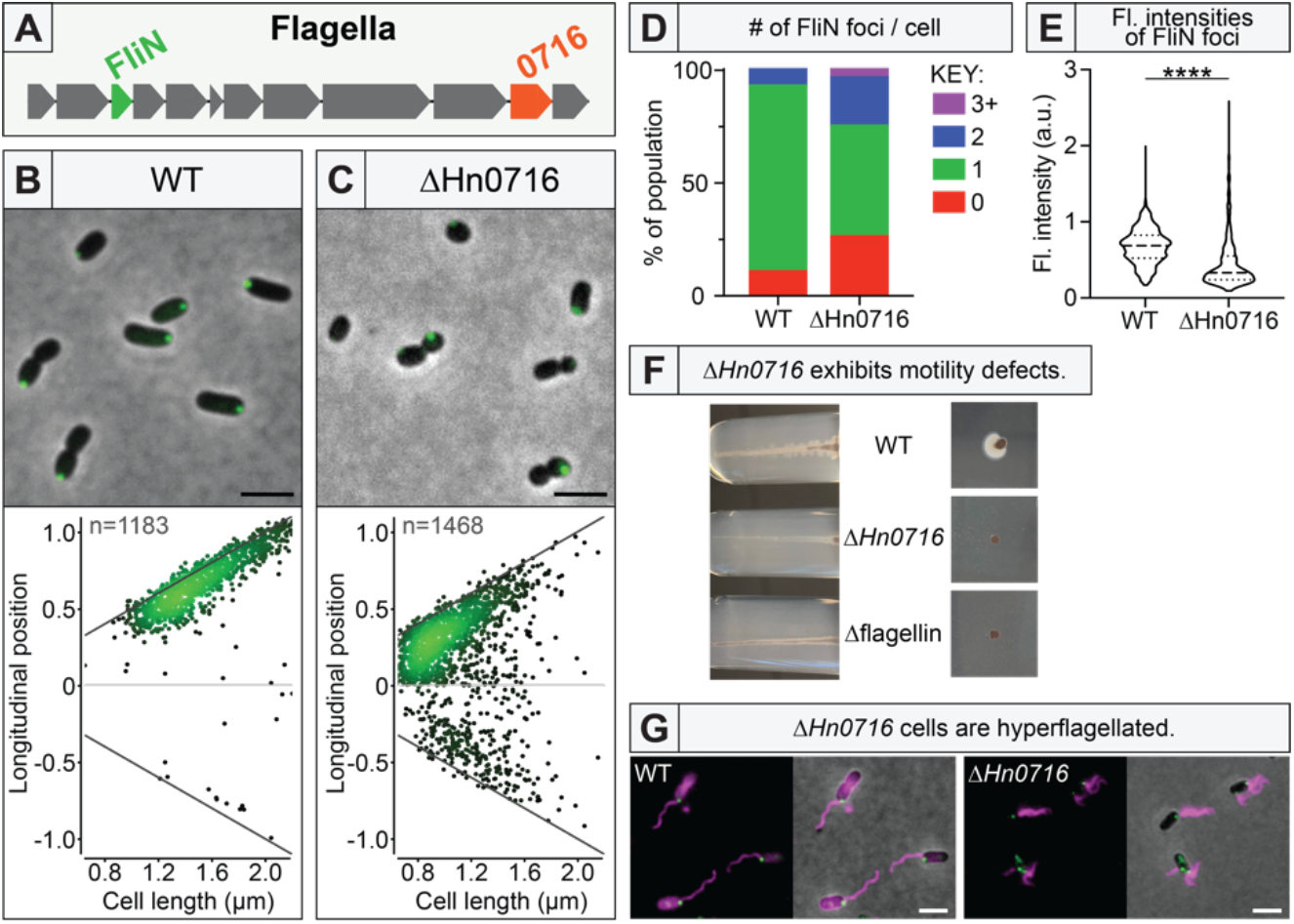
*Hn0716* is required for regulating flagella position and copy number. **(A)** *Hn0716* is found near flagella-associated genes, suggesting it is involved in positioning flagella. **(B)** Flagella localization was visualized by labelling a component of the cytoplasmic ring of the flagellar basal body (*fliN*) with mNeonGreen to form mNG-FliN. Population analysis of foci localization: Cells were analyzed and quantified using MicrobeJ. On the x-axis, cells are organized according to increasing cell length. The foci on the graphs represent where the foci are found along the length of the cell. The pole of the cell that is closest to the FliN focus is oriented to the top. Foci under the mid-cell mark represent a second focus in the cell. WT cells typically had a single polar FliN focus. Scale bar: 2 μm **(C)** In Δ*Hn0716* cells, FliN foci were more randomly positioned along the cell length. Scale bar: 2 μm **(D)** 85% of WT cells had a single focus whereas only 45% of Δ*Hn0716* cells had a single focus. Also, instead of having a single polar focus, mutant cells were 3.5 times more likely to have two or more foci and 2.5 times more likely to have no foci at all. **(E)** When FliN foci were present in Δ*Hn0716* cells, the foci were much dimmer than those of WT cells. Wilcoxon test p-value < 0.0001 **(F)** Δ*Hn0716* was not motile in motility assays. **(G)** WT cells had a single polar flagellum next to the FliN focus. Δ*Hn0716* cells typically had multiple flagella, often as tufts. Scale bar: 2 μm

**Figure S6:**
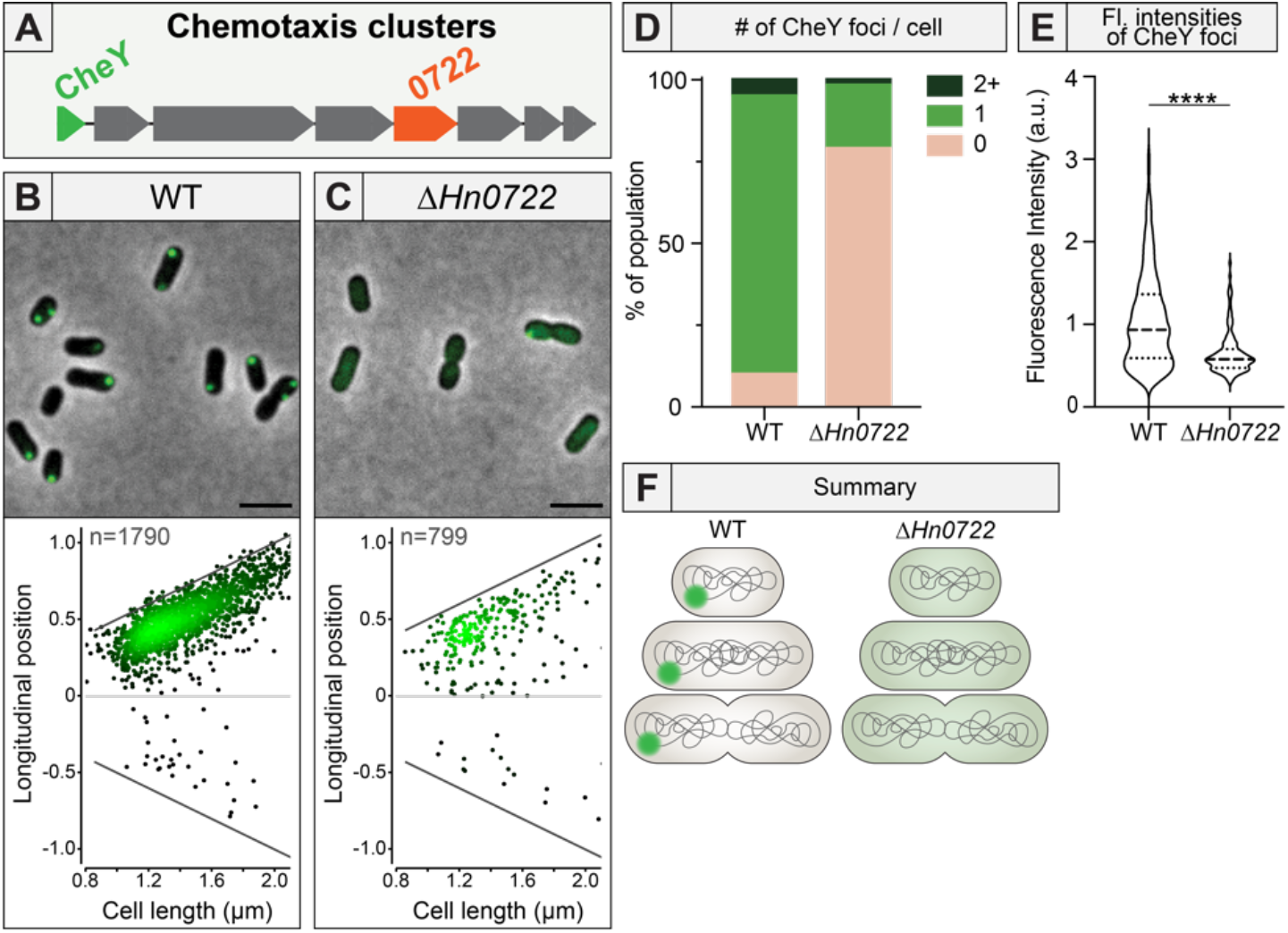
*Hn0722* is required for chemotaxis cluster positioning. **(A)** *Hn0722* is found near chemotaxis-associated genes, suggesting its involvement in positioning chemotaxis clusters. **(B)** Chemotaxis clusters were visualized by labelling the response regulator (*cheY*) with mNeonGreen to form CheY-mNG. Population analysis of foci localization: Cells were analyzed and quantified using MicrobeJ. On the x-axis, cells are organized according to increasing cell length. The foci on the graphs represent where the foci are found along the length of the cell. The pole of the cell that is closest to the CheY focus is oriented to the top. Foci under the mid-cell mark represent a second focus in the cell. WT cells had a single CheY polar focus. Scale bar: 2 μm **(C)** Δ*Hn0722* mutant cells typically had no CheY foci. Scale bar: 2 μm **(D)** 87% of WT cells had one focus whereas only 19% of Δ*Hn0722* cells had a single focus. **(E)** When Δ*Hn0722* cells did have a detectable focus, the foci were much dimmer in fluorescence intensity compared to WT. Wilcoxon test p-value < 0.0001 **(F)** *Hn0722* is critical for chemotaxis cluster assembly and positioning in *H. neapolitanus*.

**Figure S7:**
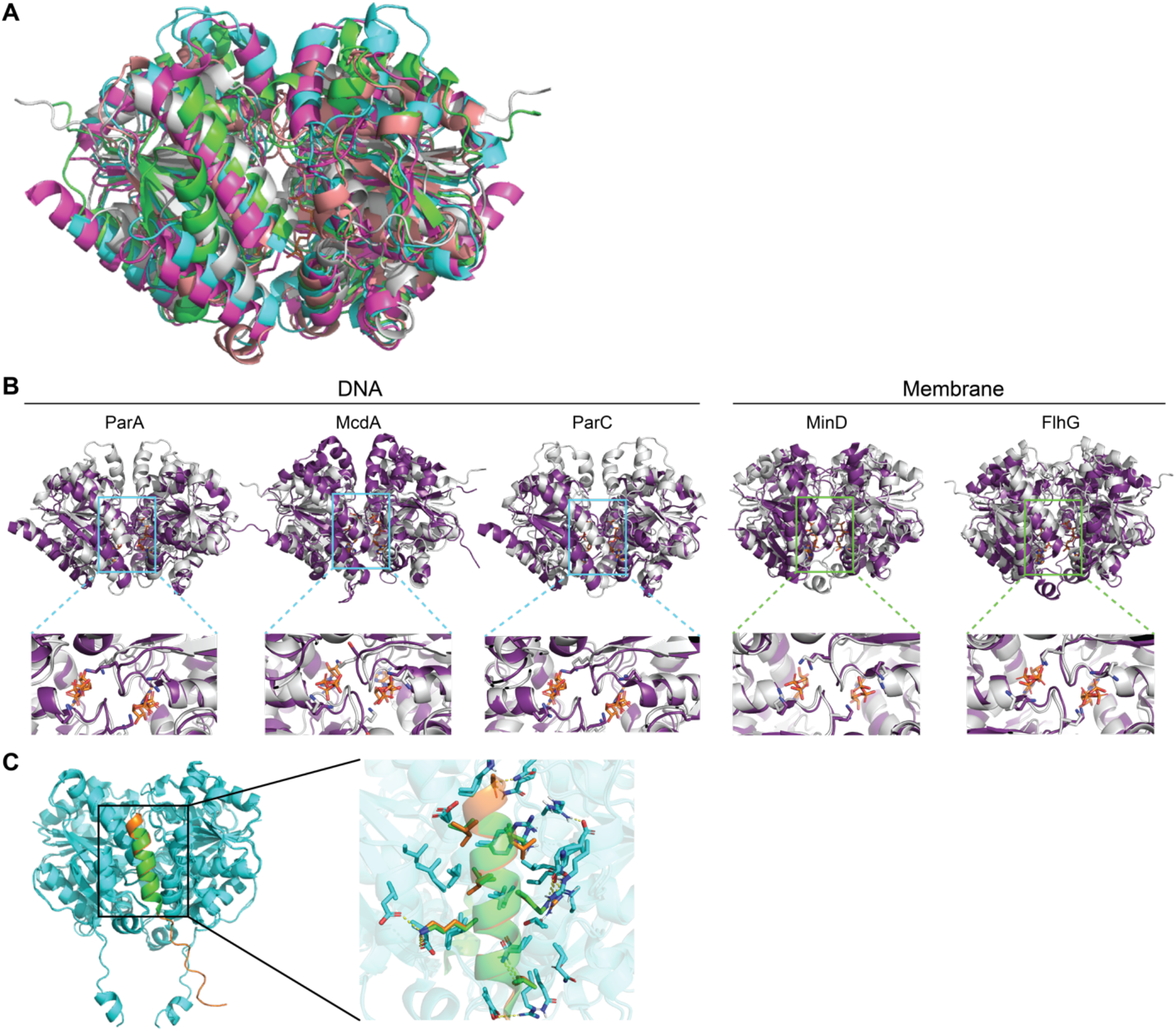
AlphaFold2 structural predictions for the A/D ATPases of *H. neapolitanus*. **(A)** AlphaFold2 predicted models of A/D ATPases in *H. neapolitanus* overlaid showing similar sandwich homodimers. ParA (magenta), McdA (gray), ParC (cyan), MinD (salmon), and FlhG (green). **(B)** Predicted A/D ATPase structures from *H. neapolitanus* (purple) overlaid with experimentally determined homologs (gray) from the Protein Data Bank (PDB). ParA, McdA, ParC, MinD, and FlhG were overlaid with PDB ID #’s 5U1G, 6NOP, 5U1G 3QPL, and 4R23 respectively. ATP-binding pockets are zoomed in below - the indicated signature lysine defines the A/D ATPase family and interacts with the γ-phosphate of the ATP molecule (orange) bound to the opposing monomer. **(C)** *In silico* alanine-substitution simulations identify the *H. neapolitanus* MinE residues (orange) experimentally shown to be important for docking the *Escherichia coli* MinE peptide (green) onto the MinD dimer in cyan (PDB ID 3Q9L). Predicted *H. neapolitanus* MinD dimer is shown overlaid in cyan.

## Table Legends

**Table S1:** ParA/MinD family ATPase hits across the bacterial domain.

**Table S2**: Amino acids comprising the interfaces of ParA/MinD family ATPases in *H. neapolitanus* predicted to be important specificity determinants for homodimerization (Tab 1), positioning matrix association (Tab 2), and partner protein binding (Tab 3). Tab 4 shows *in silico* alanine-substitution mutagenesis and the resulting ΔΔG values for all residues comprising the N-terminal peptides of partner proteins docked onto their cognate A/D ATPase.

## Movie Legends

**Movie 1: *Hn2335* is required for chromosome segregation in *H. neapolitanus*. (A)** Time-lapse microscopy of ParB-mNG foci (green) and SYTOX-stained nucleoids (magenta) showing newborn WT cells with a single ParB focus at mid-cell, which then splits into two foci that bidirectionally segregate towards the quarter positions of the growing cell. Foci positioning at the quarters of the cell was maintained, which then became the mid-cell position of each daughter cell following division. **(B)** In the Δ*Hn2335* (Δ*parA*) mutant, faithful chromosome segregation and inheritance were lost, resulting in polyploid cells that continued to divide and non-viable anucleate cells. Arrow highlights the invagination-dependent spooling of the chromosome immediately prior to complete septation. Phase Contrast (blue) shows cell perimeter. Videos accelerated ∼2,400 x real time.

**Movie 2: *Hn1364* is required for cell division positioning. (A)** Time-lapse microscopy shows WT cells dividing at mid-cell. **(B)** Δ*Hn1364* (Δ*minD*) cells divide asymmetrically. **(C)** Δ*minD* cells occasionally undergo multiple divisions simultaneously along the cell length. Phase Contrast (blue) shows cell perimeter. Arrows highlight division sites. Videos accelerated ∼2,400 x real time in (A, B) and ∼19,000 x real time in (C).

**Movie 3: Carboxysome positioning is determined by McdA, the A/D ATPase encoded in the carboxysome operon. (A)** Time-lapse microscopy of fluorescent-labelled carboxysomes (green) in dividing WT cells shows that carboxysomes are dynamically positioned along the cell length throughout the cell cycle and across multiple generations. **(B)** In Δ*Hn0912* (Δ*mcdA*) cells, carboxysome aggregates were static at the cell poles throughout the cell cycle and across multiple generations. Phase Contrast (blue) shows cell perimeter. Videos accelerated ∼2,400 x real time.

**Movie 4:** Time-lapse microscopy of ParB-mNG foci (green) in a Δ*flhG* mutant shows that chromosome segregation remains functional. But, anucleate cells form because longer cells only have a single chromosome, suggesting a defect in chromosome replication. Phase Contrast (blue) shows cell perimeter. Video accelerated ∼2,400 x real time.

**Movie 5: Carboxysomes are inherited in anucleate cells**. Time-lapse microscopy of fluorescent-labelled carboxysomes (green) in **(A)** Δ*parA*, **(B)** Δ*minD*, and **(C)** Δ*flhG* strains showing that carboxysomes can be inherited in anucleate cells. Carboxysomes in the to-be-anucleate cell bundled up immediately adjacent to the division plane (green arrows). Carboxysome bundling was coincident with chromosome extrusion through the invaginating septum just prior to complete division and asymmetric chromosome inheritance. After septation, the carboxysome bundle was explosively liberated from the anucleate cell pole, resulting in multiple freely diffusible carboxysome foci. Anucleate cells harboring carboxysomes did not divide further. Phase Contrast (blue) shows cell perimeter. Videos accelerated ∼2,400 x real time.

## References

Atmakuri, K., Cascales, E., Burton, O.T., Banta, L.M., and Christie, P.J. (2007). Agrobacterium ParA/MinD-like VirC1 spatially coordinates early conjugative DNA transfer reactions. EMBO J 26, 2540–2551.

Barlow, K.A., Ó Conchúir, S., Thompson, S., Suresh, P., Lucas, J.E., Heinonen, M., and Kortemme, T. (2018). Flex ddG: Rosetta Ensemble-Based Estimation of Changes in Protein-Protein Binding Affinity upon Mutation. J. Phys. Chem. B 122, 5389–5399.

Baxter, J.C., and Funnell, B.E. (2014). Plasmid Partition Mechanisms. Microbiol. Spectr. 2.

Blagotinsek, V., Schwan, M., Steinchen, W., Mrusek, D., Hook, J.C., Rossmann, F., Freibert, S.A., Kratzat, H., Murat, G., Kressler, D., et al. (2020). An ATP-dependent partner switch links flagellar C-ring assembly with gene expression. Proc. Natl. Acad. Sci. 117, 20826–20835.

Blair, K.M., Turner, L., Winkelman, J.T., Berg, H.C., and Kearns, D.B. (2008). A molecular clutch disables flagella in the Bacillus subtilis biofilm. Science (80-.). 320, 1636–1638.

Briegel, A., Ortega, D.R., Tocheva, E.I., Wuichet, K., Li, Z., Chen, S., Müller, A., Iancu, C. V., Murphy, G.E., Dobro, M.J., et al. (2009). Universal architecture of bacterial chemoreceptor arrays. Proc. Natl. Acad. Sci. 106, 17181–17186.

Campos-García, J., Nájera, R., Camarena, L., and Soberón-Chávez, G. (2000). The Pseudomonas aeruginosa motR gene involved in regulation of bacterial motility. FEMS Microbiol. Lett. 184, 57–62.

Chang, Y., Zhang, K., Carroll, B.L., Zhao, X., Charon, N.W., Norris, S.J., Motaleb, M.A., Li, C., and Liu, J. (2020). Molecular mechanism for rotational switching of the bacterial flagellar motor. Nat. Struct. Mol. Biol. 2020 2711 27, 1041–1047.

Dunham, T.D., Xu, W., Funnell, B.E., and Schumacher, M.A. (2009). Structural basis for ADP-mediated transcriptional regulation by P1 and P7 ParA. EMBO J 28, 1792–1802.

Hakim, P., Hoang, Y., and Vecchiarelli, A.G. (2021). Dissection of the ATPase active site of McdA reveals the sequential steps essential for carboxysome distribution. https://Doi.Org/10.1091/Mbc.E21-03-0151 32, ar11.

Hester, C.M., and Lutkenhaus, J. (2007). Soj (ParA) DNA binding is mediated by conserved arginines and is essential for plasmid segregation. Proc Natl Acad Sci USA 104, 20326–20331.

Hu, Z.L., and Lutkenhaus, J. (2003). A conserved sequence at the C-terminus of MinD is required for binding to the membrane and targeting MinC to the septum. Mol Microbiol 47, 345–355.

Jalal, A.S.B., and Le, T.B.K. (2020). Bacterial chromosome segregation by the ParABS system. Open Biol. 10, 200097.

Jones, C.W., and Armitage, J.P. (2015). Positioning of bacterial chemoreceptors. Trends Microbiol. 23, 247–256.

Jumper, J., Evans, R., Pritzel, A., Green, T., Figurnov, M., Ronneberger, O., Tunyasuvunakool, K., Bates, R., Žídek, A., Potapenko, A., et al. (2021). Highly accurate protein structure prediction with AlphaFold. Nature 596, 583–589.

Kentner, D., and Sourjik, V. (2009). Dynamic map of protein interactions in the Escherichia coli chemotaxis pathway. Mol. Syst. Biol. 5, 238.

Kerfeld, C.A., Aussignargues, C., Zarzycki, J., Cai, F., and Sutter, M. (2018). Bacterial microcompartments. Nat. Rev. Microbiol. 16, 277–290.

Kiekebusch, D., and Thanbichler, M. (2014). Spatiotemporal organization of microbial cells by protein concentration gradients. Trends Microbiol. 22, 65–73.

Kiekebusch, D., Michie, K.A., Essen, L.-O., Löwe, J., and Thanbichler, M. (2012). Localized Dimerization and Nucleoid Binding Drive Gradient Formation by the Bacterial Cell Division Inhibitor MipZ. Mol. Cell.

Koonin, E. V (1993). A Superfamily of ATPases with Diverse Functions Containing Either Classical or Deviant ATP-binding Motif. J Mol Biol 229, 1165–1174.

Kühn, M.J., Schmidt, F.K., Farthing, N.E., Rossmann, F.M., Helm, B., Wilson, L.G., Eckhardt, B., and Thormann, K.M. (2018). Spatial arrangement of several flagellins within bacterial flagella improves motility in different environments. Nat. Commun. 9, 1–12.

Kusumoto, A., Kamisaka, K., Yakushi, T., Terashima, H., Shinohara, A., and Homma, M. (2006). Regulation of polar flagellar number by the flhF and flhG genes in Vibrio alginolyticus. J. Biochem. 139, 113–121.

Leipe, D.D., Wolf, Y.I., Koonin, E. V., and Aravind, L. (2002). Classification and evolution of P-loop GTPases and related ATPases. J. Mol. Biol. 317, 41–72.

Leonard, T.A., Butler, P.J., and Löwe, J. (2005). Bacterial chromosome segregation: Structure and DNA binding of the Soj dimer - A conserved biological switch. EMBO J.

Li, S.C., and Ng, Y.K. (2010). Calibur: A tool for clustering large numbers of protein decoys. BMC Bioinformatics 11.

Livny, J., Yamaichi, Y., and Waldor, M.K. (2007). Distribution of centromere-like parS sites in bacteria: Insights from comparative genomics. J. Bacteriol. 189, 8693–8703.

Lutkenhaus, J. (2007). Assembly Dynamics of the Bacterial MinCDE System and Spatial Regulation of the Z Ring. Annu. Rev. Biochem. 76, 539–562.

Lutkenhaus, J. (2012). The ParA/MinD family puts things in their place. Trends Microbiol. 20, 411–418.

Maccready, J.S., and Vecchiarelli, A.G. (2021). Positioning the model bacterial organelle, the carboxysome. MBio 12.

MacCready, J.S., Hakim, P., Young, E.J., Hu, L., Liu, J., Osteryoung, K.W., Vecchiarelli, A.G., and Ducat, D.C. (2018). Protein gradients on the nucleoid position the carbon-fixing organelles of cyanobacteria. Elife 7.

MacCready, J.S., Basalla, J.L., and Vecchiarelli, A.G. (2020). Origin and Evolution of Carboxysome Positioning Systems in Cyanobacteria. Mol. Biol. Evol. 37, 1434–1451.

MacCready, J.S., Tran, L., Basalla, J.L., Hakim, P., and Vecchiarelli, A.G. (2021). The McdAB system positions α-carboxysomes in proteobacteria. Mol. Microbiol. 00, 1–21.

Mirdita, M., Schütze, K., Moriwaki, Y., Heo, L., Ovchinnikov, S., and Steinegger, M. (2022). ColabFold - Making protein folding accessible to all. BioRxiv 2021.08.15.456425.

Nußbaum, P., Ithurbide, S., Walsh, J.C., Patro, M., Delpech, F., Rodriguez-Franco, M., Curmi, P.M.G., Duggin, I.G., Quax, T.E.F., and Albers, S.V. (2020). An Oscillating MinD Protein Determines the Cellular Positioning of the Motility Machinery in Archaea. Curr. Biol. 30, 4956–4972.e4.

Park, K.-T., Wu, W., Battaile, K.P., Lovell, S., Holyoak, T., and Lutkenhaus, J. (2011). The Min Oscillator Uses MinD-Dependent Conformational Changes in MinE to Spatially Regulate Cytokinesis. Cell 146, 396–407.

Park, K.-T., Wu, W., Lovell, S., and Lutkenhaus, J. (2012). Mechanism of the asymmetric activation of the MinD ATPase by MinE. Mol Microbiol 85, 271–281.

Raskin, D.M., and De Boer, P.A.J. (1999). Rapid pole-to-pole oscillation of a protein required for directing division to the middle of Escherichia coli. Proc. Natl. Acad. Sci. U. S. A.

Raveh, B., London, N., Zimmerman, L., and Schueler-Furman, O. (2011). Rosetta FlexPepDockab-initio: Simultaneous folding, docking and refinement of peptides onto their receptors. PLoS One 6.

Rillema, R., MacCready, J.S., and Vecchiarelli, A.G. (2020). Cyanobacterial growth and morphology are influenced by carboxysome positioning and temperature. BioRxiv 2020.06.01.127845.

Ringgaard, S., Schirner, K., Davis, B.M., and Waldor, M.K. (2011). A family of ParA-like ATPases promotes cell pole maturation by facilitating polar localization of chemotaxis proteins. Genes Dev 25, 1544–1555.

Ringgaard, S., Zepeda-Rivera, M., Wu, X., Schirner, K., Davis, B.M., and Waldor, M.K. (2014). ParP prevents dissociation of CheA from chemotactic signaling arrays and tethers them to a polar anchor. Proc. Natl. Acad. Sci. U. S. A. 111.

Roberts, M.A.J., Wadhams, G.H., Hadfield, K.A., Tickner, S., and Armitage, J.P. (2012). ParA-like protein uses nonspecific chromosomal DNA binding to partition protein complexes. Proc. Natl. Acad. Sci. U. S. A. 109, 6698–6703.

Saha, C.K., Sanches Pires, R., Brolin, H., Delannoy, M., and Atkinson, G.C. (2021). FlaGs and webFlaGs: discovering novel biology through the analysis of gene neighbourhood conservation. Bioinformatics 37, 1312–1314.

Savage, D.F., Afonso, B., Chen, A.H., and Silver, P.A. (2010). Spatially Ordered Dynamics of the Bacterial Carbon Fixation Machinery. Science (80-.). 327, 1258–1261.

Schuhmacher, J.S., Rossmann, F., Dempwolff, F., Knauer, C., Altegoer, F., Steinchen, W., Dörrich, A.K., Klingl, A., Stephan, M., Linne, U., et al. (2015a). MinD-like ATPase FlhG effects location and number of bacterial flagella during C-ring assembly. Proc. Natl. Acad. Sci. U. S. A. 112, 3092–3097.

Schuhmacher, J.S., Thormann, K.M., and Bange, G. (2015b). How bacteria maintain location and number of flagella? FEMS Microbiol. Rev. 39, 812–822.

Schumacher, D., Harms, A., Bergeler, S., Frey, E., and Søgaard-Andersen, L. (2021). Pomx, a para/mind atpase activating protein, is a triple regulator of cell division in myxococcus xanthus. Elife 10.

Schumacher, M.A., Ye, Q., Barge, M.T., Zampini, M., BarillÃ, D., and Hayes, F. (2012). Structural Mechanism of ATP-induced Polymerization of the Partition Factor ParF. J Biol Chem 287, 26146–26154.

Schwille, P., and Diez, S. (2009). Synthetic biology of minimal systems. https://Doi.Org/10.1080/10409230903074549 44, 223–242.

Shan, S. (2016). ATPase and GTPase Tangos Drive Intracellular Protein Transport. Trends Biochem. Sci. 41, 1050–1060.

Sourjik, V., and Berg, H.C. (2000). Localization of components of the chemotaxis machinery of Escherichia coli using fluorescent protein fusions. Mol. Microbiol. 37, 740–751.

Szeto, T.H., Rowland, S.L., Rothfield, L.I., and King, G.F. (2002). Membrane localization of MinD is mediated by a C-terminal motif that is conserved across eubacteria, archaea, and chloroplasts. Proc Natl Acad Sci USA 99, 15693–15698.

Thompson, S.R., Wadhams, G.H., and Armitage, J.P. (2006). The positioning of cytoplasmic protein clusters in bacteria. Proc. Natl. Acad. Sci. U. S. A. 103, 8209–8214.

Tilly, K., Checroun, C., and Rosa, P.A. (2012). Requirements for Borrelia burgdorferi plasmid maintenance. Plasmid 68, 1–12.

Turmo, A., Gonzalez-Esquer, C.R., and Kerfeld, C.A. (2017). Carboxysomes: metabolic modules for CO_2_ fixation. FEMS Microbiol. Lett. 364.

Vecchiarelli, A.G., Han, Y.W., Tan, X., Mizuuchi, M., Ghirlando, R., Biertümpfel, C., Funnell, B.E., and Mizuuchi, K. (2010). ATP control of dynamic P1 ParA-DNA interactions: A key role for the nucleoid in plasmid partition. Mol. Microbiol.

Vecchiarelli, A.G., Mizuuchi, K., and Funnell, B.E. (2012). Surfing biological surfaces: exploiting the nucleoid for partition and transport in bacteria. Mol. Microbiol. 86, 513–523.

Wu, W., Park, K.-T., Holyoak, T., and Lutkenhaus, J. (2011). Determination of the structure of the MinD-ATP complex reveals the orientation of MinD on the membrane and the relative location of the binding sites for MinE and MinC. Mol. Microbiol. 79, 1515–1528.

Yu, X., and Margolin, W. (1999). FtsZ ring clusters in min and partition mutants: role of both the Min system and the nucleoid in regulating FtsZ ring localization. Mol Microbiol 32, 315–326.

